# Mechanism-Based and Computational Modeling of Hydrogen Sulfide Biogenesis Inhibition - Interfacial Inhibition

**DOI:** 10.1101/2022.12.06.519292

**Authors:** Laurent Le Corre, Dominique Padovani

## Abstract

Hydrogen sulfide (H2S) is a gaseous signaling molecule that participates in various signaling functions in health and diseases. The tetrameric cystathionine γ-lyase (CSE) contributes to H2S biogenesis and several investigations provide evidence on the pharmacological modulation of CSE as a potential target for the treatment of a multitude of conditions. D-penicillamine (D-pen) has recently been reported to selectively impede CSE-catalyzed H_2_S production but the molecular bases for such inhibitory effect have not been investigated. In this study, we report that D-pen follows a mixed-inhibition mechanism to inhibit both cystathionine (CST) cleavage and H_2_S biogenesis by human CSE. To decipher the molecular mechanisms underlying such a mixed inhibition, we performed docking and molecular dynamics (MD) simulations. Interestingly, MD analysis of CST binding reveals a likely active site configuration prior to gem-diamine intermediate formation, particularly H-bond formation between the amino group of the substrate and the O3’ of PLP. Similar analyses realized with both CST and D-pen identified three potent interfacial ligand-binding sites for D-pen and offered a rational for D-pen effect. Thus, inhibitor binding not only induces the creation of an entirely new interacting network at the vicinity of the interface between enzyme subunits, but it also exerts long range effects by propagating to the active site. Overall, our study paved the way for the design of new allosteric interfacial inhibitory compounds that will specifically modulate H_2_S biogenesis by cystathionine γ-lyase.

**Graphical abstract:** 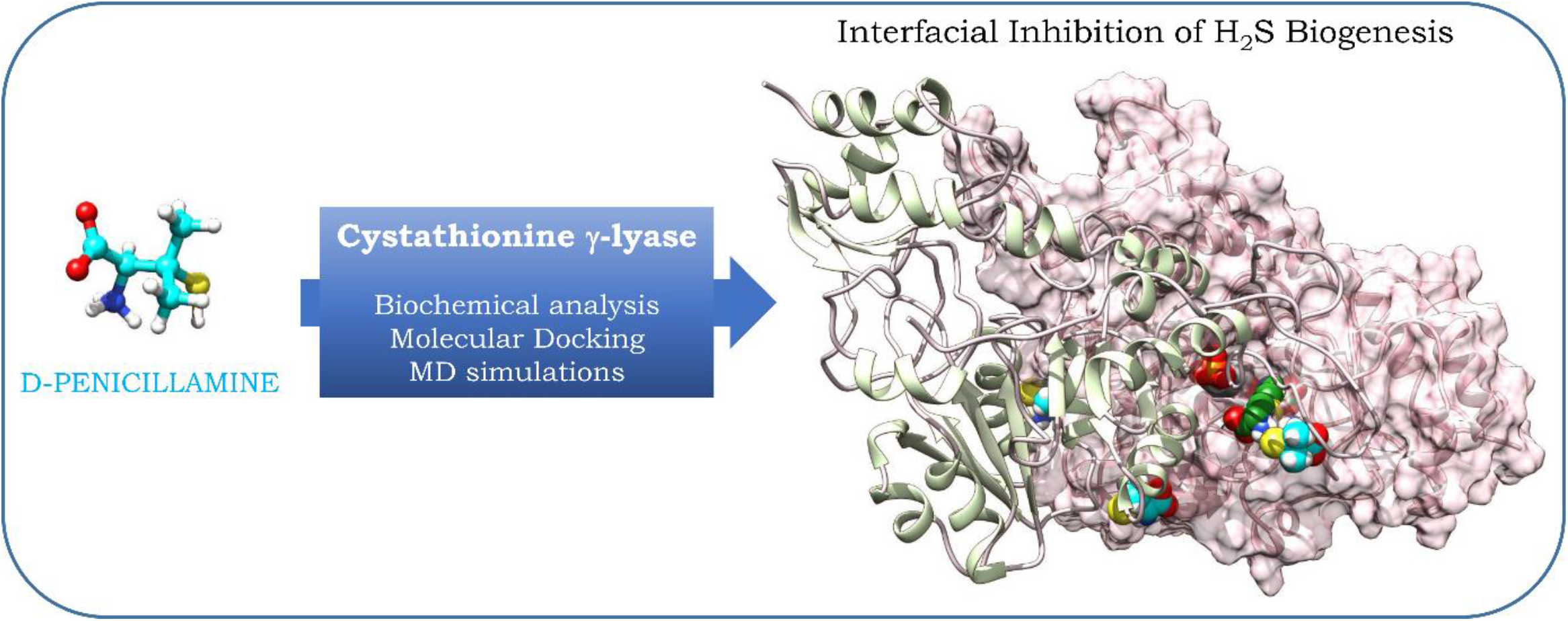

---

Hydrogen sulfide (H_2_S) is a gaseous signaling molecule that participates in various signaling functions in the vascular and neuronal systems, and in the regulation of the inflammatory response. It also contributes to the adaptive response and acts as a pleiotropic pro-resolving gaseous mediator in inflammatory diseases (1-6). In mammals, endogenous biogenesis of H_2_S is carried out by the transsulfuration pathway (TSP), constituted of two pyridoxal 5’-phosphate (PLP)-dependent enzymes, cystathionine β-synthase (CBS) and cystathionine γ-lyase (CSE), and by the mitochondrial catabolism of cysteine (Cys) that involves cysteine aminotransferase (CAT) in combination with 3-mercaptopyruvate sulfur transferase (3-MST). The TSP, with the methionine cycle to which it is connected, has also singular significance in the metabolism of sulfur compounds as well as in the maintenance of cellular homeostasis through methylation processes thanks to the control of S-adenosylmethionine (SAM) levels. In addition, the TSP represents an important source of the semi-essential amino acid Cys in mammals and its inhibition results in a net decrease of glutathione (GSH) levels in tissues (1).

In the past decades, various studies have been undertaken to decode the pivotal role of the TSP in cellular redox homeostasis in health and diseases. For example, decrypting the regulation of the TSP at the genetic level not only permitted apprehending the role of H_2_S metabolism in the modulation of inflammatory processes and the cellular response to various stress challenges, but also led to the development of sulfide-donor drugs that are currently investigated for the treatment of several diseases (2-6). Similarly, comprehending the role of the TSP in health and diseases revealed that the various systems participating in H_2_S biogenesis are up-regulated in various cancers (7, 8) and regulated upon stress challenges (4, 9). Also, H_2_S levels are modulated through up- or down-regulation of CSE in inflammatory joint disease, non-alcoholic fatty liver disease, cardiovascular or neurodegenerative diseases (10-14). Last, decoding the regulation of the TSP at the protein level revealed that both enzymes from the TSP are subjected to allosteric regulation through post-translational modifications (15-17) and that CBS heme redox status and function are regulated by gasotransmitters such as nitric oxide or carbon monoxide (1). Interestingly, these last observations recently permitted to understand how to circumvent chemotherapeutic drug resistance in breast and ovarian cancer cells (18,19).

Therefore, in a context in which the systems participating in H_2_S metabolism are regulated upon stress challenges, the pharmacological modulation of H_2_S metabolism could be investigated for the treatment of various diseases. In addition, due to the importance of CSE in H_2_S biogenesis, compounds modulating its activity seem essential for investigating and deciphering its contribution to cell physiology, the adaptive response and the modulation of diseases. However, few inhibitors of CSE have been identified so far, and many of these compounds display significant drawbacks such as lack of selectivity for CSE over CBS due to the PLP-dependent activity of both enzyme or even lack of studies to decipher their selectivity against other PLP-dependent enzymes (20-22). Hence, Cystathionine γ-lyase is a tetrameric enzyme that belongs to the γ-family of PLP-dependent enzymes, and catalyzes Cys or H_2_S biogenesis from cystathionine or Cys and/or homocysteine, respectively (23, 24) (**Fig. 1**). In the resting state, the PLP cofactor forms an internal lysine-derived aldimine with Lys212 and is further stabilized in the active site through several interactions with surrounding amino acid residues, *e*.*g*. conservation of the N1-pyridine of the PLP in a protonated state through H-bonding with the side-chain of Glu187 that is maintained in the proper orientation through interacting with Thr189, π- π stacking of the pyridine ring of PLP with Tyr114, H-bonding of the O3’ atom from PLP with the side-chain of Asn161 and the Nε-H+ of Lys212, H-bonding of the _3_ OPO ^2-^ group with the backbone of Leu91 and Gly90, the side-chains of Ser209, Thr211, as well as the side-chains of *Tyr60 and *Arg62 from the adjacent subunit (**Fig. 1*B***). During catalysis, the amino group of the substrate will replace Lys212 through a transimination reaction to form an external aldimine intermediate that will further participate in α,β- or α,γ-elimination depending on the substrate (**Scheme 1**) (24, 25).

**Figure 1.**
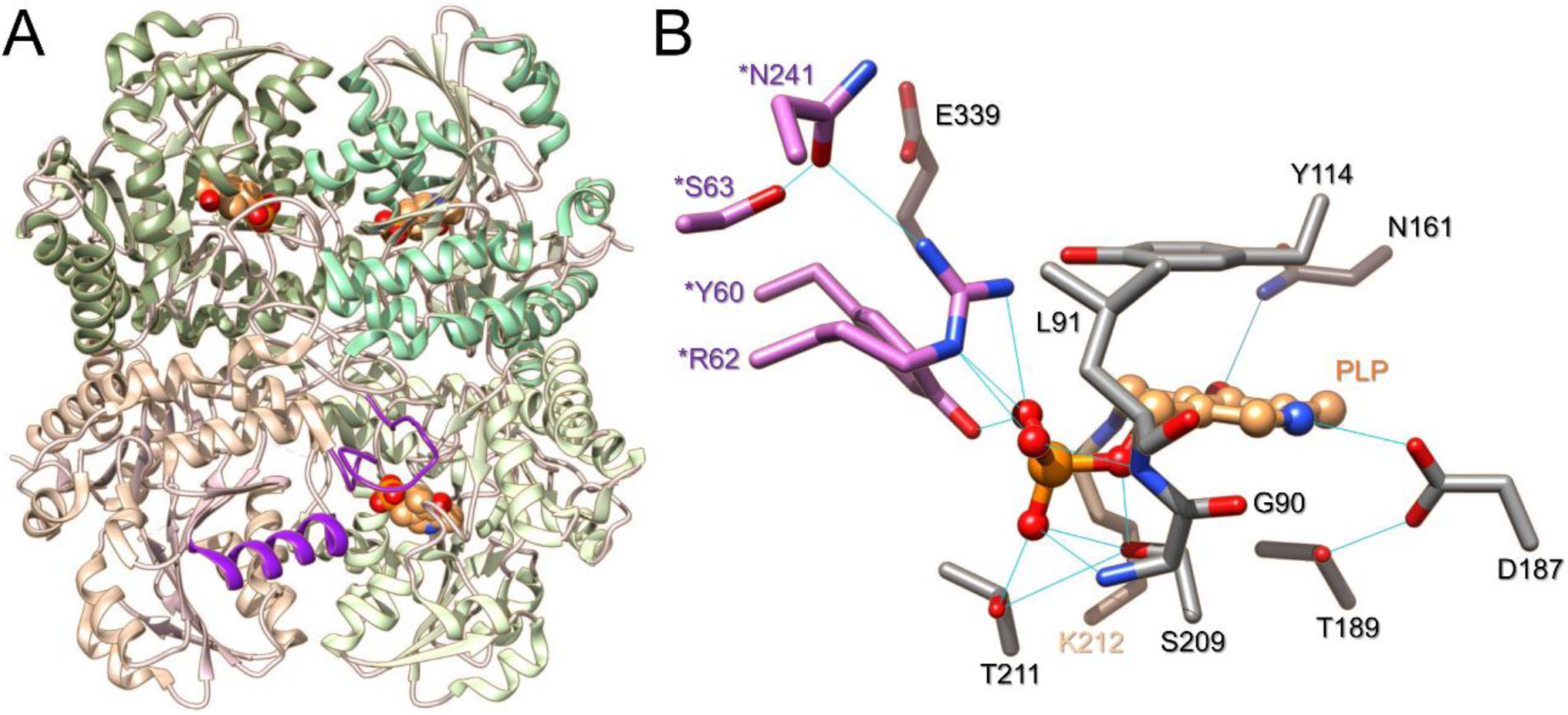
Structure of Cystathionine γ-lyase. (**A**) Structure of CSE showing its tetrameric organization (PDB code: 2NMP). Each subunit of CSE is color-coded differently. The PLP cofactor is shown in brown. Structural elements from the adjacent B subunit (loop44-65 and α-helix229-244) contributing residues to the active site of the A subunit are color-coded in purple. (**B**) Close-up of CSE active site from the A subunit. Important amino acid residues are represented in stick representation. The PLP cofactor (brown) establishes an imine bond at its C4’ position with Lys212 (tan color) and is stabilized through various hydrogen bonds (represented in light blue lines) with surrounding amino acid residues, including residues from the adjacent B subunit (colored in orchid).

**Scheme 1.**
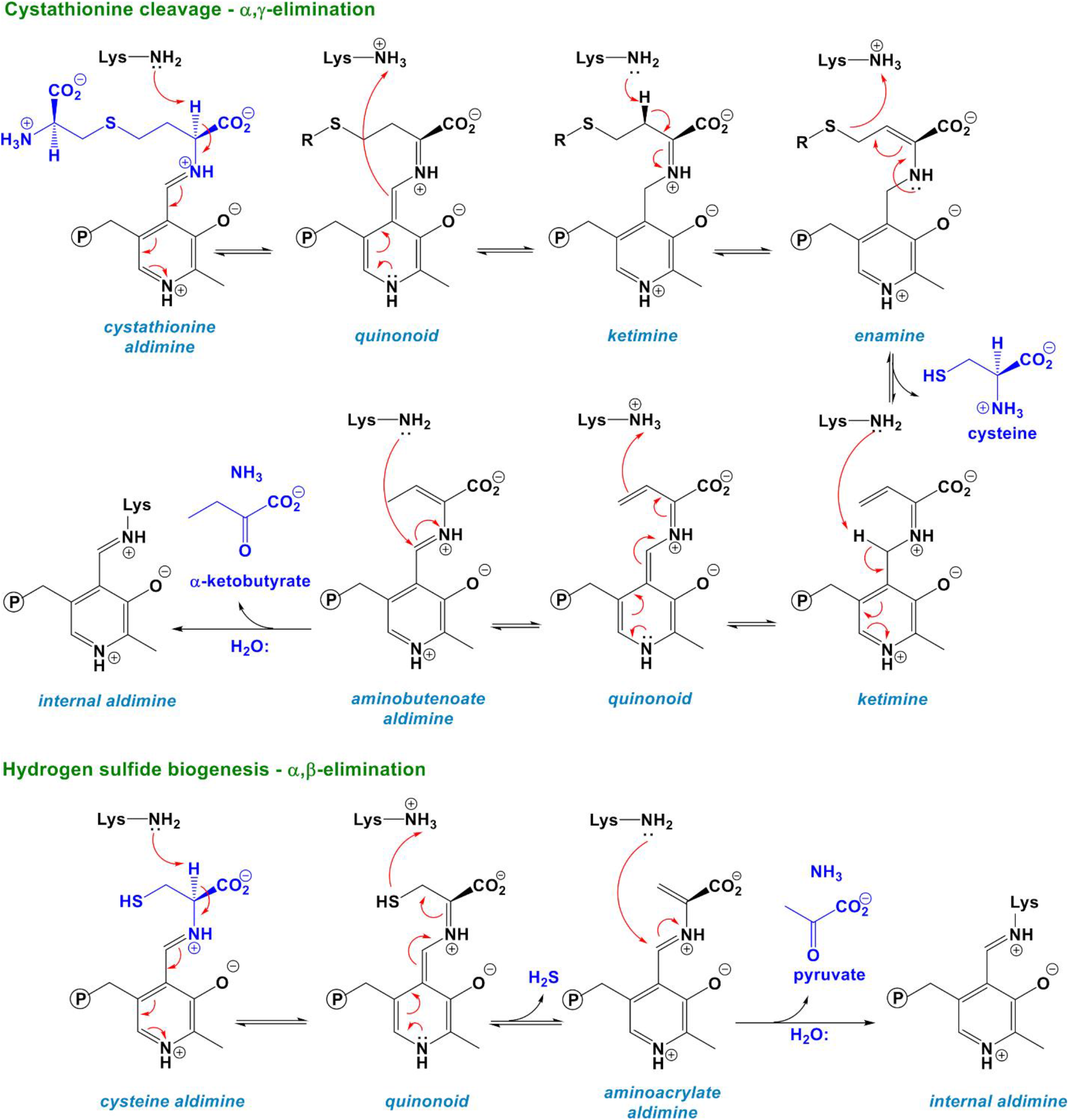
Main reactions catalyzed by cystathionine γ-lyase. The α,γ-elimination of cystathionine leads to cysteine formation (top) while the α,β-elimination of cysteine produces the biological gaseous mediator H2S (bottom). The steps leading from the internal aldimine to the formation of the external aldimine are not shown for reasons of clarity.

The scarcity of selective CSE inhibitors has prompted recent interest in this area with high-throughput (virtual) screening (HTS) of compound libraries that identified iminoquinolinone derivative, N-propargyl D-cysteine derivative, aurintricarboxylic acid, isoxsuprine and 2-arylidene hydrazinecarbodithioates as selective modulators of CSE activity (26-30). In addition, a semi-rational *in silico* drug screening study combined with chemical synthesis (31) and a screening of substrate analogs (32) reported *(R)*-2-oxo-*N*-(prop-2-yn-1-yl)thiazolidine-4-carboxamide and S-3-carboxypropyl-L-cysteine as potent selective inhibitors of CSE, with the later inducing stable aminoacrylate formation. Finally, D-penicillamine (D-pen), a drug already used in the management of rheumatoid arthritis and Wilson’s disease, recently repurposed in cancer treatment and responsible for the reversible inhibition of catalase, has also been reported to selectively inhibit CSE but the molecular bases for such inhibitory effect have not been investigated (33-35).

In this study, we report a combined kinetic, molecular docking and molecular dynamics analysis of D-penicillamine capacity to modulate the activity of CSE. Our kinetic study reveals that D-Pen inhibits both the cystathionine and cysteine cleavage reactions catalyzed by CSE through a mixed inhibition mechanism. Our molecular dynamics analysis of cystathionine binding shows that the substrate adopts a specific active configuration prior to gem-diamine intermediate formation by establishing a strong interacting network with surrounding amino acid residues and by forming H-bond contact with the O3’ atom of PLP. Our studies show that this competent state is no longer visible in the presence of D-penicillamine, which binds to the interface of two adjacent subunits. Interestingly, inhibitor binding not only induces the creation of an entirely new interacting network at the vicinity of the interface between enzyme subunits, but also exerts long range effects by propagating to the active site thus explaining its particular mode of inhibition.

## Experimental procedures

### Materials

All chemicals were purchased from Sigma-Aldrich and used as-is. L-(+)-cystathionine was purchased from Bertin Technologies or Interchim Inc.

### Expression and purification of recombinant proteins

The plasmid containing the human CSE gene (pET-28-based expression vector incorporating a tobacco etch virus (TEV)-cleavable N-terminal His tag fusion) was a kind gift from Tobias Karlberg (Structural Genomics Consortium, Karolinska Institute, Sweden) (36). The plasmid was transformed into One Shot BL-21 Star (DE3) *E. coli* (Invitrogen) and grown overnight in 3L of a Luria Bertani (LB) medium supplemented with kanamycin (60 mg/L). The next morning, expression of hCSE was induced by the addition of 0.5 mM isopropyl β-d-1-thiogalactopyranoside (IPTG), and the culture was grown for an additionnal 20-24 hours at 20 °C. Cells were harvested by centrifugation (4,000 rpm, 15 min) and the cell pellet was resuspendended then sonicated 10 min (20 cycles; 30 s on, 2 min off) in 50 mM sodium phosphate buffer, pH 8.0, 300 mM NaCl (buffer A) containing 20 mM imidazole, 1 mM dithiothreitol and EDTA-free protease inhibitor (Roche Applied Science). The solution was then centrifuged for 1 h at 20,000 rpm and 4 °C. The supernatant was filtered through a MF-Millipore Membrane Filter (0.45 µm pore size) then loaded on a 5 mL His-Trap HP column (GE Healthcare) pre-equilibrated with buffer A and CSE was eluted with a gradient from 20 mM to 250 mM imidazole in buffer A. The fractions of interest, as judged by 12 % SDS-PAGE, were pooled, concentrated with an Amicon Ultra-15 centrifugal filter unit (30 kDa cut-off) and dialyzed against 100 mM HEPES, pH 7.4, 150 mM NaCl (buffer B). The protein was then loaded onto a HiLoad 16/600 Superdex 200 (GE Healthcare) and eluted at 0.5 mL/min with buffer B. The fractions of interest were pooled, washed with 100 mM HEPES at pH 7.4 and concentrated as above, and then stored at -80 °C. Noteworthy, the activity assays were performed with a tag-less CSE. To remove the N-terminal 6xHis-tag, the CSE protein obtained after affinity purification was dialysed against buffer A, and then incubated overnight at 4°C with His-tagged TEV protease at a molar ratio CSE:TEV of 20:1. The mixture was then passed through a 5 mL His-Trap FF column (GE Healthcare) equilibrated with buffer A. Then, the flow-through containing CSE was subjected to dialysis and gel filtration as decribed above.

The plamid pKR793-TEV coding for TEV containing a 6xHis-tag was a kind gift of Ludovic Pecqueur (Collège de France, France). It was transformed into BL21-CodonPlus (DE3)-RIL competent cells (Agilent Technologies) and cells were grown overnight at 37 °C in LB medium supplemented with ampicilline (100 mg/L) and chloramphenicol (15 mg/L). Then, 1L of LB medium complemented with the same antibiotics was ensemenced at 2% with the preculture. Cells were grown to OD_600_ of 0.8 at 37 °C. TEV overexpression was induced by adding 0.5 mM IPTG, and growth was continued at 25 °C overnight. Cells were harvested as decribed above and the cell pellet was treated as previously described for hCSE. Sonication was then performed for 10 min. (10 cycles; 1 min on, 3 min off). After sonication and centrifugation, TEV was purified via the batch method using Ni-NTA Superflow resin (Qiagen), as described by the manufacturer. TEV was concentrated as above, and then stored at -80 °C. The concentration of CSE and TEV was determined using the Bradford reagent (Bio-Rad) with bovine serum albumin as a standard.

### Enzymatic assay to detect cysteine production from cystathionine

Usually, the formation of cysteine from cystathionine is measured in a continuous DTNB (5,5’-dithiobis-2-nitrobenzoic acid) activity assay (24). However, due to the presence of D-penicillamine in the reaction mixtures, we were not able to use such assay. Instead, we followed the formation of α-ketobutyrate, another reaction product formed during the α,γ-cleavage of cystathionine by CSE, through a discontinuous 2,4-dinitrophenylhydrazine (DNPH) activity assay. Briefly, 500 μL of the assay mixture containing varying concentrations of cystathionine (0-3 mM) and D-penicillamine (up to 20 mM) were preincubated for 5 min in 100 mM HEPES buffer at pH 7.4 and 37 °C. The reaction was initiated by adding 2-5 μg of CSE and at the desired time points (2-5 min), 100 μL aliquots of the reaction mixture were quenched by adding 10 % v/v of 10% trichloroacetic acid. Samples were centrifuged (13,000 rpm, 10 min at 4 °C) and the supernatant was mixed with 20 μl of a DNPH solution (2 mM in 2 M HCl). After 10 min incubation at 37°C, the reaction mixture was loaded on a 96-well microplate and mixed with 280 μL of 1 M NaOH. Appropriate control experiments lacking substrate or enzyme were performed in parallel. After 10 min, the absorbance at 430 nm was recorded using a BioTek PowerWave XS microplate reader. The concentration of α-ketobutyrate in the reaction mixture was calculated using a standard curve generated with known concentrations of α-ketobutyrate. The effect of D-penicillamine on CSE was determined by estimating *K*_M_^app^ and *V*_max_^app^ values for CSE-catalyzed α-ketobutyrate production from cystathionine (CST) a each D-penicillamine (D-pen) concentration, and using Equation 1.

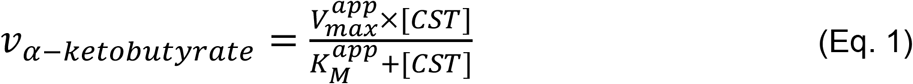

Then, the CSE-inhibitor dissociation constants *K*_ic_ for competitive inhibition and *K*_iu_ for uncompetitive inhibition were calculated by plotting *V*_*max*_^app^/*K*_*M*_^app^ or *V*_*max*_^app^ as a function of D-penicillamine concentration, respectively, and using Equations 2 or 3 (37).

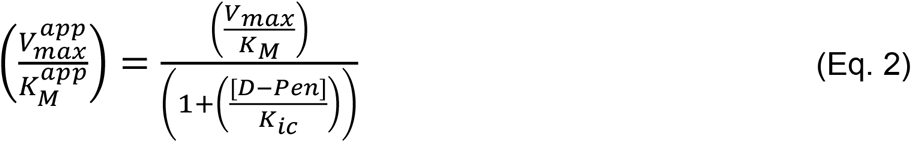

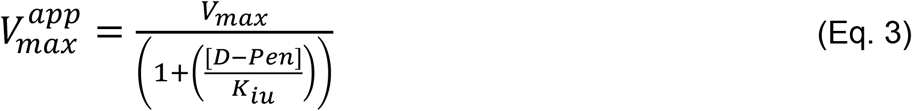

### Enzymatic assay to measure H_2_S production from cysteine

Production of H_2_S from cysteine by CSE was measured spectrophotometrically by the lead acetate assay as previously described (24). First, the effect of D-penicillamine on the activity of CSE was determined from a reaction mixture containing 1 mM cysteine, 0.4 mM lead acetate and varying concentration of D-penicillamine (up to 100 mM) in 200 mM HEPES at pH 7.4. After 3 min of preincubation of the assay mixture at 37 °C, the reaction was initiated by adding 40 μg of CSE. Then, the formation of lead sulfide resulting from the reactivity of H_2_S with lead acetate was continuously monitored at 390 nm. The half inhibitory concentration (IC_50_) value was obtained by plotting the relative activity of CSE (A, in per cent) as a function of the concentration of D-pen and by fitting the data with Equation 4, where ΔA is the maximal effect of D-pen on the activity of CSE and n is the Hillslope that characterizes the slope of the curve at its midpoint.

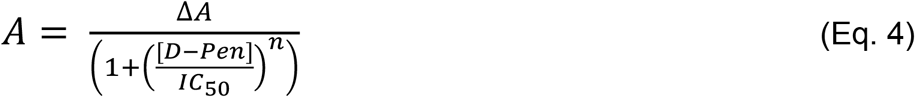

Second, the effect of D-penicillamine on the kinetic parameters of CSE-catalyzed H_2_S generation was estimated in the same activity assay. Briefly, the reaction mixture containing varying concentrations of cysteine (1-10 mM), lead acetate (0.4 mM) and D-penicillamine (0, 1, 5 or 10 mM) was preincubated for 3 min in 100 mM HEPES buffer at pH 7.4 and 37 °C. Next, 40 μg of CSE was added to the assay mixture to initiate the reaction, which was followed at 390 nm. The molar extinction coefficient for lead sulfide previously determined was then used to obtain *v*_H2S_ (24). The CSE-D-penicillamine dissociation constants for competitive and uncompetitive inhibition were calculated by plotting *v*_H2S_ as a function of cysteine concentration and using Equations 5.

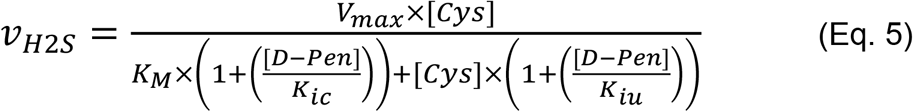

### Fluorescence spectroscopy

D-penicillamine binding experiments were performed by following PLP fluorescence (λ_exc_ = 423 nm; λ_em_^max^ = 495 nm) on a fluorimeter (Shimadzu) equiped with a thermostating device. Tag-less CSE (1 μM) was incubated in 100 mM HEPES at pH 7.4 and 20 °C with successive aliquots (0.5-2 μL) of D-penicillamine stock solutions prepared in the same buffer. Fluorescence emission spectra were recorded 5 min after D-pen addition. Equilibrium dissociation constant (*K*_d_) of D-pen to CSE was determined by plotting the relative decrease in fluorescence emission at 495 nm against the D-penicillamine concentration and by using equation 6, whit F_0_ the minimal relative fluorescence at saturating D-pen concentration, ΔF_max_ the maximal relative decrease in fluoresence emission provoked by D-pen binding to CSE, and n the Hill coefficient that leasures the potential cooperativity in ligand binding process.

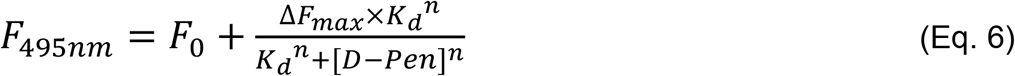

### Molecular docking and dynamics studies

The crystal structure of the PLP bound human CSE tetrameric complex was downloaded from the Protein Data Bank (PDB ID: 2NMP, https://www.pdb.org). A dimeric structure of CSE formed by chains A and B was prepared using the Prepare Protein module in Biovia Discovery Studio® (DS) 2016 with the default parameters and the CHARMM force field. Briefly, all crystallographic water molecules, except HOH 512, were removed. Bond orders were assigned, hydrogen and missing atoms were added. The protonation states on protein and PLP were adjusted at pH 7.4. The imine bond between Lys212 and C4’ atom of PLP was created using the Build and Edit Protein tool of DS. Finally, the protein/PLP complex 3D structure was minimized using the Adopted Basis Newton-Raphson (ABNR) algorithm.

The 3D-coordinates of L-(+)-cystathionine and D-penicillamine were generated using the Prepare Ligand module in DS with ionization states calculated at pH 7.4. Random conformations of each ligand (8 for D-penicillamin and 26 for cystathionine) were generated for docking experiments using the BEST algorithm (38) to improve the coverage of the conformational space. Docking studies were performed using CDOCKER (39) implemented in DS 2016 and all parameters were kept at their default values. Potential binding sites in the CSE protein structure were determined using the receptor cavities tool of DS and the fpocket application (40) within the Galaxy server (https://cheminformatics.usegalaxy.eu/). Pose clustering, based on the RMSD values, was performed using the RMSD tool of DS. Best poses within each cluster, based on the CDOCKER Interaction Energy and CDOCKER Energy scores, were further refined through the In Situ Ligand Minimization protocol of DS. Following this, the binding mode of compounds was thoroughly examined using the Analysis Ligand Poses module of DS.

To assess the structural stability of the L-(+)-cystathionine- and D-penicillamine-bound CSE, 50 ns of molecular dynamics simulations were run starting from best docking models using the CHARMM36m force field and the NAMD protocol (41) implemented in DS 2021. Protein-ligand complexes were solvated in an orthorhombic box using a TIP3P water model. Periodic boundary conditions were applied with a minimum distance of 7 Å from periodic boundary and Na+Cl-counter ions were added to neutralize the system. Solvated complexes were subjected to 50000 steps of energy minimization with the ABNR algorithm followed by 20 ps of heating from 50 to 300K at constant volume, restraining the position of protein with a force constant of 10 kcal/mol Å, then equilibrated during 100 ps in the NPT ensemble (300K, 1 atm) without any constraints. Finally, MD simulations were performed in the NPT ensemble (300K, 1 atm). Langevin Dynamics- and Langevin Piston methods were applied to control the temperature and the pressure. Short-range electrostatic and Van der Waals interactions were computed with a 12 Å cut-off distance and long-range electrostatic interactions were treated by the Particle Mesh Ewald (PME) method. All bonds with hydrogen atoms were held rigid using the SETTLE algorithm. RMSD, RMSF values, and distance calculations were generated using the Analysis Trajectory tool of DS.

## Results and discussion

### Purification and Characterization of CSE

Human recombinant CSE was purified as described previously (24) with the following modification: after Ni-NTA affinity chromatography, the protein was either treated with TEV protease to remove the N-terminal 6×His-tag or dialysed and then loaded onto a HiLoad 16/600 Superdex 200 column. The Q-Sepharose step was thus skipped. The purified full-length and tag-less proteins were judged to be pure ≥ 90-95% by gel electrophoresis (**Fig. S1**). In addition, both full-length (42) and tag-less CSE purified as a homotetramer with estimated apparent molecular masses of 152 ± 3 and 148 ± 4 kDa, respectively. Last, the absorption spectrum of purified proteins exhibits a maximum at 428 nm that is characteristic of an internal aldimine-containing CSE (24) (**Fig. S1**).

The goal of the present study was to estimate the effect of the thiol D-penicillamine on the kinetic parameters of CSE-catalyzed reactions. However, the influence of D-pen on the kinetic parameters associated with the α,γ-elimination reaction of cystathionine by human CSE could not be determined by the standard *in vitro* assay that uses Ellman’s reagent (43). Instead, we monitored the formation of α-ketobutyrate, another reaction product of the α,γ-cleavage of cystathionine (**Scheme 1**), using a discontinuous 2,4-dinitrophenylhydrazine (DNPH) activity assay. In such conditions, purified full-lenth CSE displays *V*_max_ of 2.47 ± 0.05 U/mg and *K*_M_ of 0.27 ± 0.02 mM with cystathionine as a substrate (**Fig. S2**). These values are similar to the ones determined in the classical assay (V_max_ = 2.5-3.1 U/mg and *K*_M_ = 0.28-0.40 mM) (24, 42). Tag-less CSE exhibits a *V*_max_ of 2.33 ± 0.02 U/mg and *K*_M_ of 0.25 ± 0.01 mM for the the α,γ-cleavage of cystathionine in the DNPH activity assay (**Fig. S2**). Thus, the catalytic efficiency of human CSE is not affected by removal of the N-terminal 6×His-tag. The tag-less protein was used to perform the following studies.

### Interaction of D-penicillamine with human CSE

A previous study reported that D-pen-dependent inhibition of CSE was reversed in a concentration-dependent manner by the addition of exogenous PLP (34). It was thus suggested that D-pen might exert its inhibitory effect on H_2_S biogenesis by interfering with the PLP cofactor. As a result, we investigated this likely interaction by fluorescence spectroscopy. Addition of D-penicillamine to human CSE results in a decrease in fluorescence emission of the PLP cofactor at 495 nm (**Fig. 2**), thus suggesting that D-pen binding to CSE affects the environment of this intrisic probe. Quantification of the dissociation constant between tag-less CSE and D-pen gives a *K*_d_ value of 15.0 ± 4.6 mM (n=3 ± SD), suggesting a loose interaction between both molecules.

**Figure 2.**
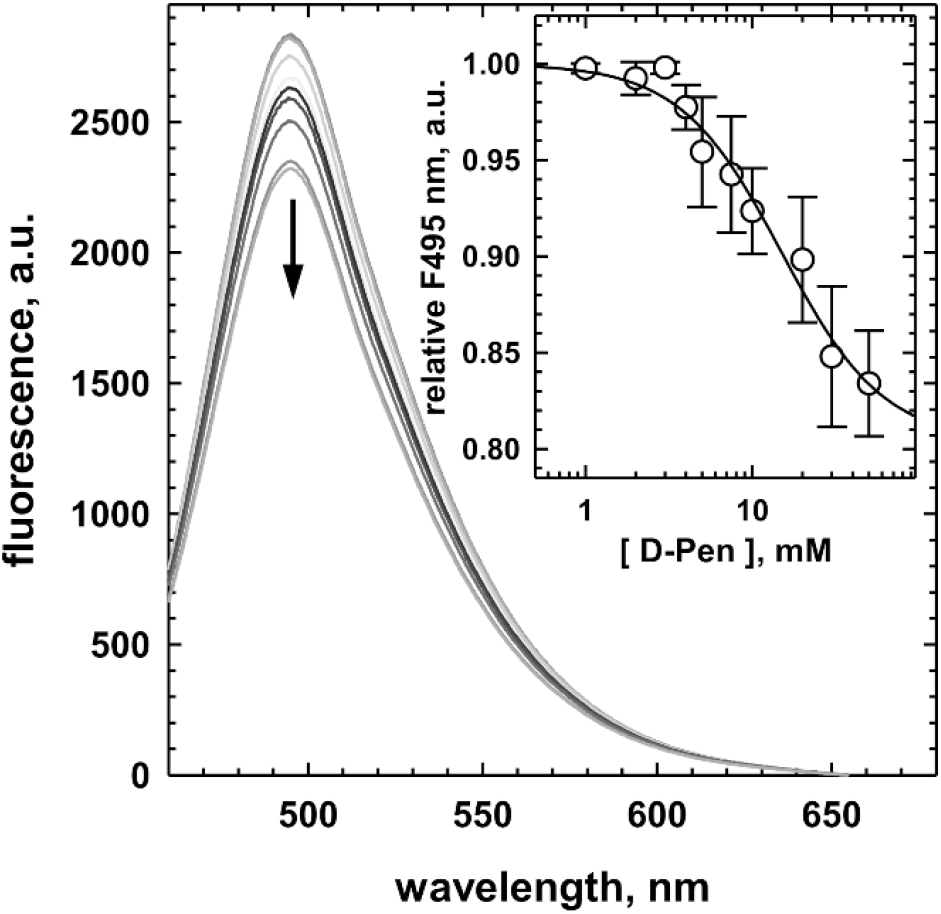
Binding of D-penicillamine to cystathionine γ-lyase. Representative experiment showing the fluorescence emission spectra (λ_exc_ = 423 nm) of recombinant human CSE (1 μM) recorded in 100 mM HEPES buffer pH 7.4 at 20 °C in the presence of increasing concentrations of D-penicillamine (up to 50 mM). The addition of D-pen induced a quenching of fluorescence. Inset: Dependence of the relative fluorescence recorded at 495 nm for D-penicillamine binding on human CSE (n=3±SD). The black line represents the fit to the experimental data with a four-parameter logistic equation to obtain the dissociation binding constant of D-pen for CSE.

### Mechanism based inhibition of D-penicillamine

Then, we studied the mode of inhibition of CSE by D-penicillamine by starting with its impact on CSE-catalyzed H_2_S biogenesis from cysteine. In the lead acetate assay, D-penicillamine exhibits an IC_50_ value of 2.04 ± 0.13 mM towards H_2_S production from 1 mM Cys (**Fig. 3*A***). This result is in stark contrast with the IC_50_ value of 0.27 mM previously measured using the same substrate concentration in the methylene blue assay (34). In this preceding study, D-pen was pre-incubated for 10 min with CSE prior to Cys addition, in contrast to the experiments described herein. The difference between both studies could thus come from a slow-binding inhibition process in which the enzyme-inhibitor complex is assembled on a time scale of several minutes or from an induced fit mechanism of inhibition by D-penicillamine (37). Accordingly, we followed the inhibition potency of D-penicillamine as a function of the pre-incubating time. Nevertheless, the inhibitory capacity of D-pen is not affected by a pre-incubation period, suggesting that D-penicillamine is not a slow-binding inhibitor. The dissimilarity between both *in vitro* studies could therefore result in part from the differences in assay conditions and from the purity of the enzyme used. Thus, Brancaleone *et al*. used the methylene blue assay which displays major drawbacks (44, 45) and they used a GST-CSE enzyme purified with a single chromatographic step (GST HiTrap FF affinity column) that leads to a 50-60 % pure enzyme (22).

**Figure 3.**
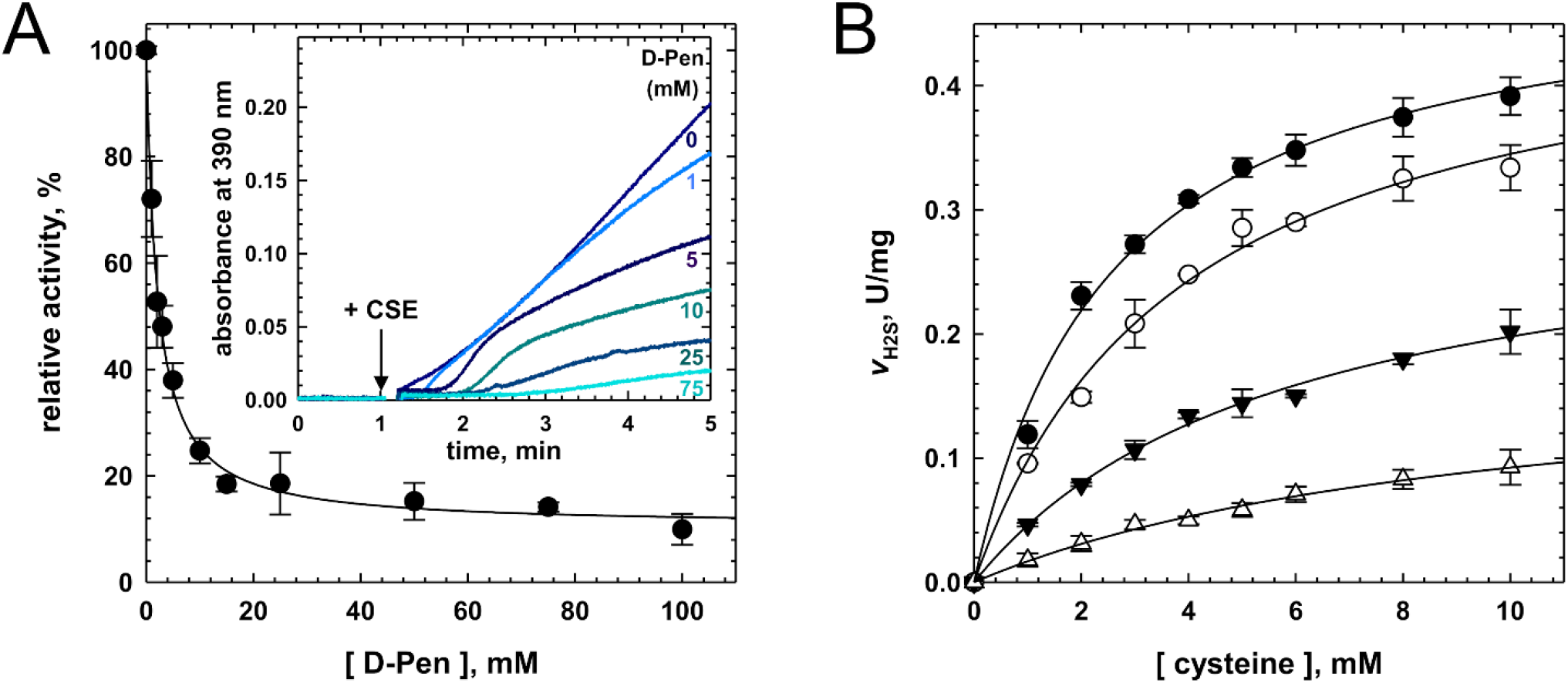
D-penicillamine inhibits hydrogen sulfide biosynthesis. (**A**) Experiments showing the dependency of H_2_S production from Cys (1 mM) by human CSE on the concentration of D-pen (up to 100 mM) in 200 mM HEPES at pH 7.4 and 37 °C. The plot (n=2±SD) was fitted with a four-parameter logistic equation and the relative IC_50_ value is reported in the text. Inset: Representative experiments showing the inhibition of human CSE activity by D-penicillamine in the H_2_S assay. The reaction mixture contained cysteine (1 mM), lead acetate (0.4 mM) and varying concentrations of D-pen (indicated on the plot) in 200 mM HEPES, pH 7.4 at 37 °C. H_2_S production was initiated by adding human CSE (40 μg) and recorded at 390 nm (formation of a PbS precipitate). (**B**) Michaelis-Menten plot analysis of CSE inhibition by D-penicillamine. Inhibition of H_2_S synthesis was determined in assay mixtures containing 1-10 mM cysteine, 0.4 mM lead acetate, CSE (40 μg) and varying concentrations of D-pen (⬤, 0 mM; ○, 1 mM; ▼, 5 mM; △, 10 mM) in 100 mM HEPES at pH 7.4 and 37 °C. The data (n ≥ 2 ± SD) were fitted as described in “Experimental Procedures” to extract the CSE-inhibitor competitive (*K*_ic_) and uncompetitive (*K*_iu_) dissociation constants.

It must be noticed that Brancaleone *et al*. observed a significant difference in the IC_50_ values exhibited by D-pen *in vitro* (IC_50_ of 0.27 mM against recombinant CSE) and *ex cellulo* (IC_50_ of 0.044 mM in homogenated mouse aorta samples). Whereas an induced fit inhibition mechanism could have explained such a variance, it could also result in part from the *in cellulo* metabolization of D-Pen into a more potent inhibitory metabolite. For example, D-pen is a substrate for D-amino acid oxidase (DAO) (46) which could thus generate dimethyl mercaptopyruvate from D-pen, the former being a substrate analog for 3-MST. Thus, either dimethyl mercaptopyruvate or D-pen itself could inhibit 3-MST, the enzyme primary responsible for H_2_S biogenesis in rodent coronary artery homogenates (47), which is under consideration. This could also explain the results obtained *ex vivo* in the presence of propargylglycine since this non-specific inhibitor of PLP-dependent enzymes will deprive 3-MST of its substrate *via* the inhibition of cysteine aminotransferase, the enzyme involved in the biogenesis of 3-mercaptopyruvate. Last, D-pen can lead to the formation of hydrogen peroxide (H_2_O_2_) through the oxidase activity of certain hemoproteins (35). H_2_O_2_ being involved in the regulation of rodent coronary artery blood flow (48), the effect produced by H_2_O_2_ thus generated could be added to the points already developed to partly explain the vasodilatation variation observed by Brancaleone *et al*. in the presence of D-pen.

Next the *K*_i_ of D-pen for CSE was assessed in the H_2_S assay in the presence of varying concentration of Cys. Surprisingly, Lineweaver-Burk analysis of the data suggests that D-pen followed a mixed inhibition mechanism (**Scheme S1**) to impede H_2_S biogenesis, since the plots intersect outside the x- and y-axes (**Fig. S3*A***). As a result, Michaelis-Menten plots were individually fitted with Equation 5 to extract the dissociation constants for competitive (*K*_*ic*_) and uncompetitive (*K*_*iu*_) inhibition, which yield values of 1.82 ± 0.35 mM and 8.60 ± 1.22 mM for *K*_*ic*_ and *K*_*iu*_, respectively (**Fig. 3*B***). Noteworthy, kinetic parameters for H_2_S biogenesis from Cys determined during these analyses (*K*_M_ = 3.03 ± 0.44 mM and V_max_ = 0.49 ± 0.03 U/mg, n=3 ± SD) are like the ones previously reported (24).

Last, CSE inhibitor could harbour different inhibition constants depending on the length of the substrate, as noted with S-3-carboxypropyl-L-cysteine with regard to both cystathionine and cysteine cleavage inhibition (32). We thus determined CSE-D-pen dissociation constants for competitive and uncompetitive inhibition during the CSE-catalyzed α,γ-elimination reaction of cystathionine (**Fig. 4**). D-pen also follows a mixed inhibition mechanism to hinder cystathionine cleavage, as suggested by Lineweaver-Burk analysis of the data that shows that the plots intersect outside the x- and y-axes (**Fig. S3*B***), with values of 2.54 ± 0.65 mM and 8.55 ± 0.89 mM for *K*_*ic*_ and *K*_*iu*_, respectively (**Fig. 4*B***). D-pen is therefore slightly less capable to competitively inhibit the cystathionine cleavage reaction than the cysteine cleavage reaction. In contrast, the dissociation constant for uncompetitive inhibition is not affected by the length of the substrate. Kinetic parameters for cystathionine cleavage determined during these analyses (*K*_M_ = 0.21 ± 0.02 mM and V_max_ = 2.27 ± 0.06 U/mg, **Fig. 4*B***) are also comparable to the ones reported above (**Fig. S2*B***).

**Figure 4.**
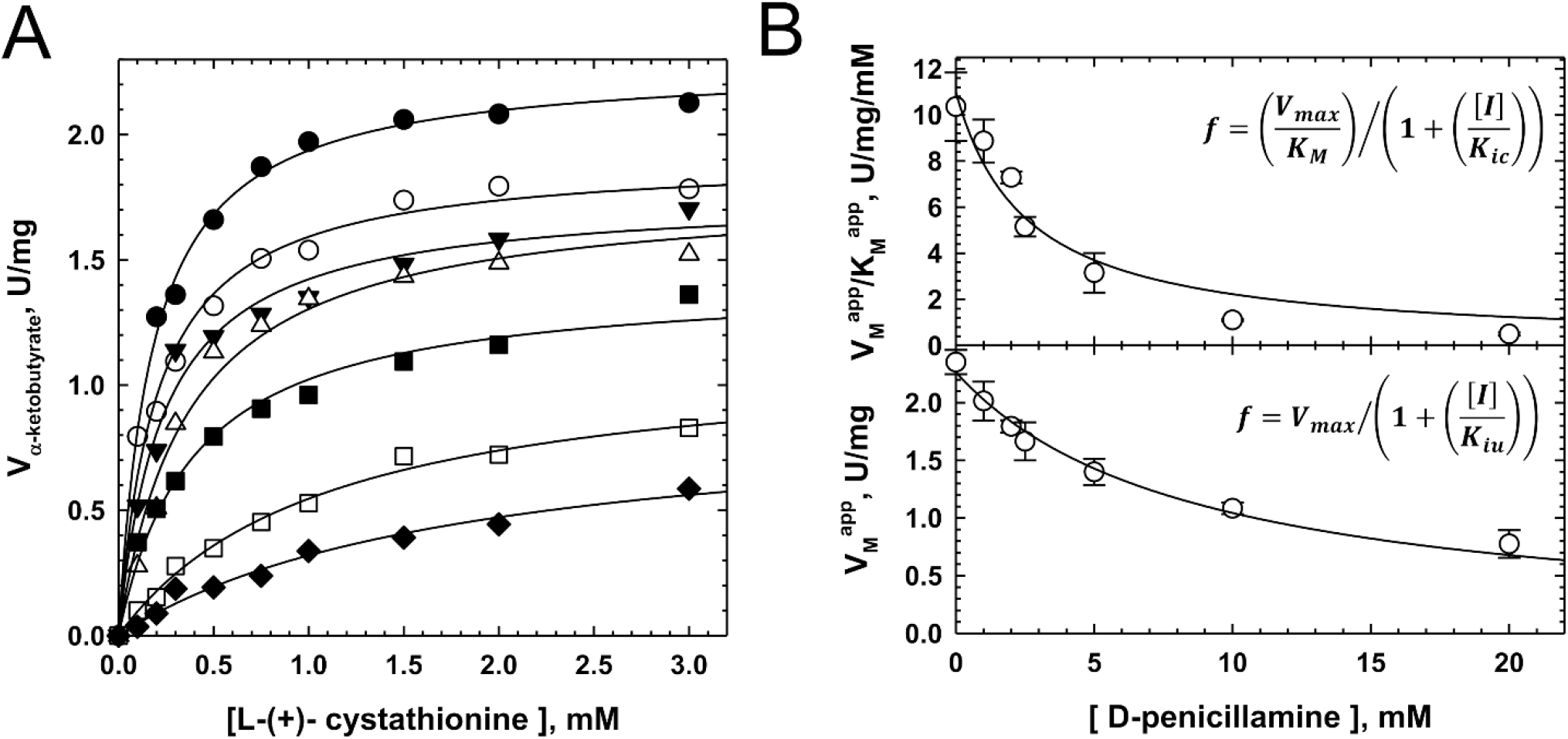
Mixed inhibition of cysteine biogenesis from cystathionine by D-penicillamine. (**A**) Representative Michaelis-Menten plot analysis of CSE inhibition by D-penicillamine. Inhibition of cysteine synthesis from L-(+)-cystathionine was determined by measuring -ketobutyrate production at 37 °C from assay mixtures containing 0.1-3 mM L-(+)-cystathionine, varying concentrations of D-pen (0-20 mM, top to bottom) and CSE (5 g) in 100 mM HEPES at pH 7.4. The data were fitted with a hyperbolic function to extract *V*_M_^app^ and *K*_M_^app^ at each D-penicillamine concentration. (**B**) Determination of the CSE-inhibitor dissociation constants *K*_ic_ for competitive inhibition (top) and *K*_iu_ for uncompetitive inhibition (bottom) (n ≥ 2 ± SD).

Altogether, our results suggest than D-pen is a mixed inhibitor of cystathionine γ-lyase (**Scheme S1**). Interestingly, despite its with weak potency to impair both cystathionine and cysteine cleavage activity of CSE, the inhibitory mode adopted by D-pen is quite peculiar as it differs from the ones reported for trifluoroalanine (43), propargylglycine (36, 43) and S-3-carboxypropyl-L-cysteine (32) that are suicide inhibitors. It also contrasts with the inhibitory mode detected for aminoethoxyvinyglycine, a slow and tight binding inhibitor of CSE (43), and the mode of action of I157172 (29) and aurintricarboxylic acid (30) or *(R)*-2-oxo-*N*-(prop-2-yn-1-yl)thiazolidine-4-carboxamide (31), respectively competitive cofactor and substrate inhibitors.

### A minimized molecular model for cystathionine γ-lyase

Accordingly, to further investigate the molecular mechanisms responsible for the unusual mixed inhibition mechanism displayed by D-penicillamine, we performed computational modeling studies. At first, we generated a structural model of CSE based on the crystal structure of CSE (PDB code: 2 NMP). Our minimized molecular model mainly differs from the crystal structure in the following points: (i) since the quaternary structure of the enzyme consists of a dimer of dimer (36), we choose to work with a minimalist model constituted of a single dimer comprising the A and B subunits; (ii) hydrogen atoms were added such as the pyridine moiety from the cofactor and the imine bond between the Nε-lysine (Lys212) and the C4’ atom of PLP are accurately protonated; (iii) the length of the aforementioned imine bond is of 1.30 Å in our model while it exhibits an unusual length of 1.96 Å in the crystal structure (PDB code: 2 NMP); (iv) last, Asn161 is no longer in H-bond contact with the O3’ atom of PLP.

To investigate the stability of our model, we then performed 50 ns molecular dynamics (MD) simulations and monitored the time evolution of the root mean square deviation (RMSD) and the per-residue root mean square fluctuation (RMSF) (**Fig. S4**). Our dimeric model is relatively stable during the time course of the simulations, and most importantly nearly all the structural elements constituting the active site fluctuate marginally during the simulations. Noteworthy, the two distinct changes in the RMSD values seen at the beginning (t = 0-0.1 ns) and at the end (t ≥ 20-30 ns) of the simulations are respectively related to the equilibration protocol and mostly to the movements of the N-terminal portion of the protein backbone that is usually implicated in the dimer-dimer interface, as shown in the per-residue RMSF (**Fig. S4**).

### Computational modeling of cystathionine binding reveals a likely active site configuration prior to gem-diamine intermediate formation

Next, we used our minimized model to perform docking and molecular dynamics (MD) simulations on cystathionine (CST) binding to the active site of CSE (**Fig. 5** and **Figs. S4-S7**). The representative docking pose from cystathionine bound to CSE (CDOCKER energy and CDOCKER interaction energy for docked cystathionine of -67.8 kcal/mol and -61.1 kcal/mol, respectively) shows that essentially four structural elements (named I to IV) contribute to the substrate access channel and the active site geometry, and also provide amino acid residues for PLP and/or substrate binding (**Figs. 5*A*** and **S5**). Thus, the cohesion and opening of the substrate access channel are mainly well-maintained thanks to the hydrogen bonding network created between the side chains of amino acid residues from structural elements I and IV (*Gln49 [an asterisk denotes an amino acid residue from the B subunit] and Glu339), from structural elements I, II and III (*Ser63, *N241, *S242 and R119), and from structural elements III and IV (Arg122, Gln123 and Asp363) (**Fig. 5*A***). Noteworthy, this network is also completed by several hydrogen bonds established between amino acid residues from the same structural element that will not be discussed in here.

**Figure 5.**
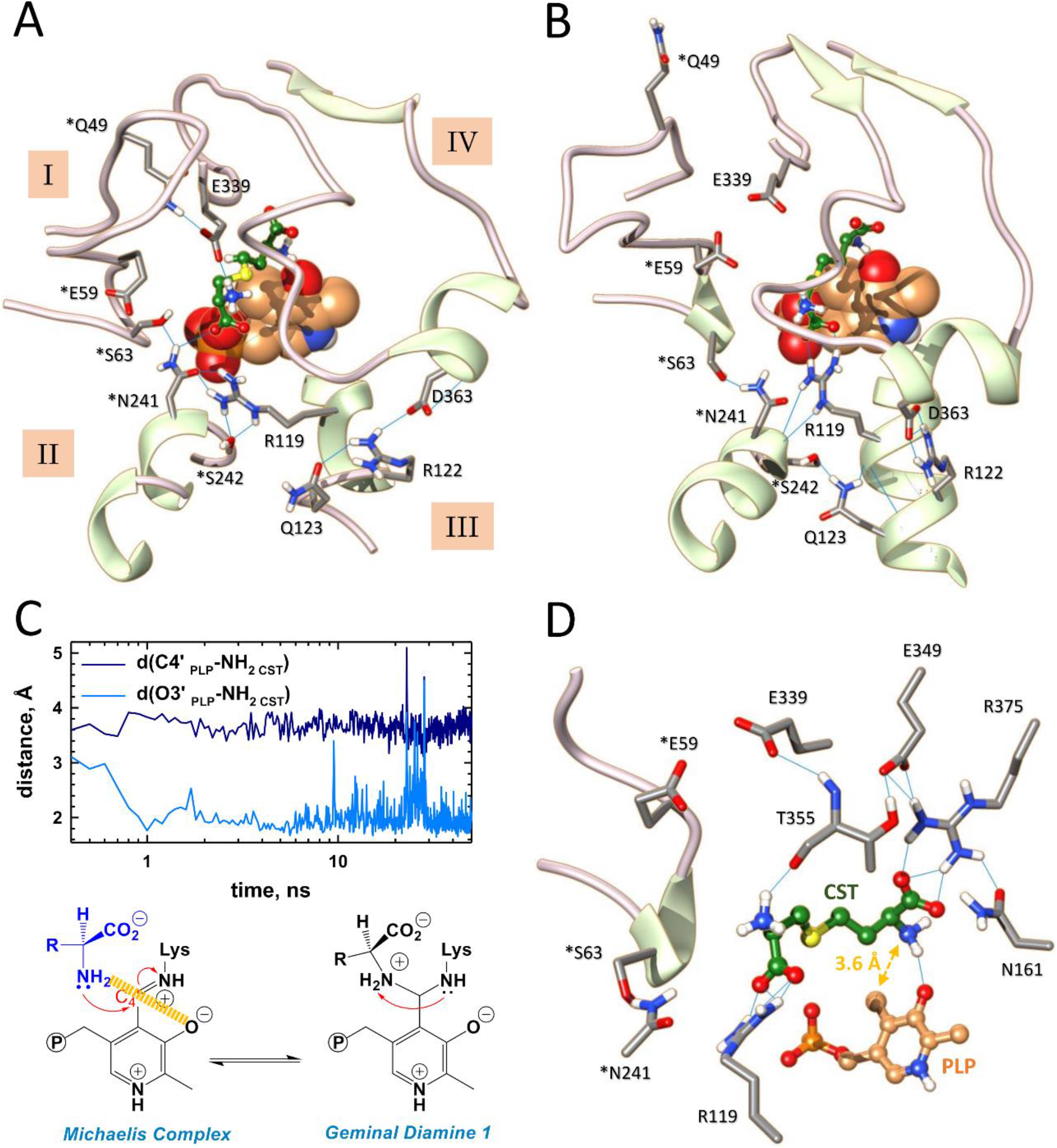
Molecular dynamics analysis of cystathionine binding reveals a likely active site configuration prior to *gem* diamine intermediate formation. (**A, B**) Structural comparison of the active site from subunit A from a representative docking pose (**A**) and a representative snapshot of MD simulation at t=50 ns (**B**). As shown, MD simulations induce structural changes as well as modifications of the hydrogen bonding network observed in the docking pose. (**C**) Plots showing average distances between the proximal nucleophilic NH_2_ of cystathionine and the C4’ (dark blue) or O3’ (light blue) atoms of PLP. (**D**) Representative snapshot of the active site from subunit A at 50 ns during MD simulation showing the interacting network of cystathionine with surrounding amino acid residues and cofactor. Particularly, an hydrogen bond is established between the proximal amino group of the substrate and the O3’ atom of PLP. *Residues from the adjacent subunit B. H-bonding are depicted by light blue lines. CST = cystathionine.

Besides, the PLP cofactor is involved into π-π stacking with Tyr114, its N1-pyridine-H^+^ forms an H-bond with Glu187, and PLP is anchored by strong hydrogen bonding between its phosphate moiety and amino acid residues from the A (Gly90, Leu91, Ser209) and B (*Arg62) subunit (**Fig. S5*A***). Additionally, the distal part of the substrate principally initiates hydrogen bond contacts at the entrance of the substrate channel with the side chains of *Arg62, *Asn241, Tyr114 that was postulated to act as a base for deprotonating the substrate (49), Glu339 that has been proposed to govern the specificity of CSE-catalyzed reactions (50, 51), and the backbone of Thr355. Last, Asn161 and Thr355 provide hydrogen-bond interactions with the proximal α-carboxyl group of cystathionine at the bottom of the substrate binding site, as does Arg375 according to previous docking experiments performed with yeast CSE (49) (**Fig. S5*B***). Interestingly, the H-bond contacts implicating Asn161 and Arg375 might be preserved along the course of CSE-catalyzed reactions as the carboxylate of an aminoacrylate bound to CSE (PDB code: 6NBA) forms hydrogen-bonding interactions with both amino acid residues (32). Also, site-directed mutagenesis experiments showed that Arg375 is important for maintaining the H_2_S-producing activity of the enzyme (51). Last, cystathionine is further stabilized by establishing van der Waals contacts with *Tyr60 and Tyr114, C-H…S and O…S interactions with the side chain of Thr355, and unconventional C-H…O and C-H…N contacts with Leu341 or Lys212, respectively.

The representative docking pose was then subjected to 50 ns molecular dynamics (MD) simulations to investigate the stability of the CSE-CST binary complex (**Fig. 5*B-D*, S4, S6*A*** and **S7*A***). While CST is stable along the entire sampling interval, its binding to CSE enhances the stability of the enzyme, as shown by the reduced RMSD (ΔRMSD = 1.67 Å over the last 20 ns) and RMSF (ΔRMSF = 0.60 Å) values observed for the CSE-CST binary complex in comparison to the ones detected for the minimized model (**Figs. S4*A* and S4*E***). MD simulations reveal changes at the vicinity of the binding site, particularly in the initial hydrogen bonding network observed in the docking pose that maintain the unity of the active site and substrate access channel (**Fig. 5*B***), in the interacting network of PLP that is now better stabilized thanks to its interaction with two new amino acid residues (*Tyr60 and Thr211) (**Fig. S6*A***) as well as in the interacting network of cystathionine (**Figs. 5*C-D*** and **S7*A***).

Importantly, the connexion between structural elements I-II, II-III and III-IV are essentially preserved despite differences with the original interacting network observed in the docking pose and regardless of structural changes in elements I-IV implicated in the substrate access channel and the active site geometry. For example, the tip of the flexible loop^44-65^ (structural element I) from the B subunit moves away from its original position in part due to the breaking of the connexion between the side chains of *Gln49 and Glu339 (**Fig. 5*B***). The later now becomes indirectly implicated in the stabilization of the substrate at the active site through H-bonding with Thr355 (**Fig. S7*A***). In addition, the guanidinium part of Arg119 flips about 90° to form a salt bridge with the distal α-carboxylate of CST while still staying connected with element II through H-bond contacts between its NH2 and Nε and the side chain of *Ser242. Also, elements II and III are connected through H-bonding between *Ser242 and Gln123, itself in H-bond contact with the backbone of Arg119 (**Fig. 5*B***). Last, element IV undergoes a rearrangement during which (i) a new β-sheet antiparallel to the existing one is formed, (ii) several residues (*e*.*g*. backbones of Ser340 and Leu341, side-chains of Thr355 and Arg375) now fall within a radius of hydrogen interaction with Glu349 localized on the newly formed anti-parallel β-sheet, (iii) Arg375 establishes a salt bridge with the proximal α-carboxylate of the substrate and hydrogen-bond contact with Asn161 (**Fig. 5*B*** and **S7*A***).

Furthermore, cystathionine is no more H-bonded to Tyr114 that establishes H-bonding contact with *Arg62 as well as S/π and O…S interactions with the substrate (**Fig. S7*A***). Noteworthy, the distance between the O atom from Tyr114 and the S atom from the substrate (2.93 Å) is inferior to the sum of the van der Waals radii of O (1.52 Å) and S (1.80 Å) (52), suggesting a stabilizing interaction between these atoms. The microenvironment created around the tyrosine hydroxyl group by the interactions described above may profoundly perturb its intrinsic p*K*_a_ thus rendering Tyr114 prone to participate into proton transfer during cystathionine cleavage, for instance into the protonation of the leaving group as suggested for CGL from *L. plantarum* (53). Furthermore, these structural changes cause the access channel from the substrate to the active site to become slightly more constrained, as indicated by the distances between the three amino acid residues *Ser63, Ala357 and Asp363 (perimeter of 38.0 Å [docking] *versus* 33.6 Å [MD]) (**table S1**). Also, cystathionine slides 1.3 Å deeper into the substrate access channel, which may facilitate its interaction with active site amino acid residues and shield the active site from solvent exposure. Altogether, these structural rearrangements and the creation of new interacting networks for the PLP and substrate possibly promote the organising of the active site prior to catalysis.

Strikingly, the monitoring of the average distances between the O3’ or C4’ atoms of PLP and the proximal substrate amino group unexpectedly reveals the formation of a H-bond contact between the O3’ atom and the amino group of cystathionine during MD simulations (**Fig. 5*C-D*** and **Table S1**). This would allow the O3’ atom to participate into substrate deprotonation though two possible mechanisms, as previously reported (54, 55). First, the O3’ atom could directly deprotonate the proximal substrate amino group as observed in L-serine hydratase from *Xanthomonas oryzae* pv. *oryzae* (54) (**Scheme S2**). Thus, the keto form of the O3’ atom (species A) would play a catalytic role during the transamination reaction by taking part in the proton transfer from the substrate to the lysine amino group implicated into Schiff base formation. The initial abstraction of a proton from the substrate amino group will generate the enol form of the O3’ atom (species B), thus disturbing the original internal H-bond between the O3’ atom and the N-H^+^ from the Schiff base (species A). This would allow the Schiff linkage to adopt a single-bond resonance form and to easily rotate around the C4’-C4(pyridine) axis, thus creating a more electrophilic carbon at the C4’ position and facilitating the nucleophilic attack of the amine group of the substrate (species C). Second, the O3’ atom could intervene in the indirect deprotonation of the substrate as suggested by QM/MM calculations during the degradation of L-methionine by methionine γ-lyase from *Pseudomonas putida* (55). In this case, the O3’ atom contributes to the water-assisted deprotonation of the substrate, which has been found more favourable and less energetic than the direct deprotonation of L-methionine by the O3’ atom to generate the nucleophilic species prior to *gem* diamine formation. A water molecule located 4.3 Å away from the O3’ atom in the resting state of CSE (PDB code: 2NMP) could accomplish this task admitting that such a mechanism is followed by CSE during the α,γ-elimination of cystathionine.

Alternate and/or complementary functions may be attributed to the formation of the H-bond contact between the O3’ atom and the nucleophilic substrate since our modeling studies have been performed with the substrate amino group already deprotonated. The H-bond linkage could first have a self-contradictory role. On the one hand it could stabilize the nucleophile, making it less reactive and thereby increasing the intrinsic barrier for the formation of the *gem* diamine intermediate. On the other hand, it could better position the nucleophilic amino group in space and therefore modulate its selectivity and favour its attack towards the C4’ atom, consequently decreasing the intrinsic barrier for the formation of the *gem* diamine (56, 57). Second, the hydrogen bond linkage could stabilize the adducts formed between PLP and the substrate, intermediates, or the product during catalysis since its occurrence has been detected in the crystal structures of the external aldimine formed between cystathionine and PLP of *L. plantarum* CGL (PDB code: 6LE4) (53) and in human CSE harbouring an aminoacrylate intermediate (PDB code: 6NBA) (32). Last, the H-bond could also modulate the electrophilic character of the C4’ atom. Thus, while the protonation state of N1-pyridine determines the absolute acidity of the Cα and C4’ of aldimines and ketimines, the protonation state of imine and O3’ modulates the relative acidities of the Cα and C4’ (58, 59). Accordingly, the snapshot captured at 50 ns during MD simulations (**Fig. 5*B*** and **5*D***) could represent an enzyme conformation needed for promoting catalysis and particularly for lowering the activation energy of the transition state leading to *gem* diamine intermediate formation.

### Prediction of D-penicillamine binding sites and docking of the ligand

Next, we performed computational studies to identify putative ligand binding sites within CSE and further carry out similar studies on the ternary complex CSE-substrate-ligand. The prediction of the ligand binding sites was performed with the Galaxy site program using the crystal structure of CSE (PDB code: 2NMP, A and B chains) and with the Cavity search program from Biovia Discovery Studio (DS) 2021 using our minimized molecular model as input. Molecular docking was performed with CDOCKER software implemented in DS. Best binding site were retained based on docking score (CDOCKER interaction energy and CDOCKER energy scores). Thus, amongst the seventy-one putative ligand-binding sites, eighteen of them were selected for docking studies based on their size as well as their location on the enzyme. Finally, three potential ligand binding sites in which D-pen binds with similar docking score were selected (**Fig. 6**). Interestingly, the 3 binding sites are all interfacial ligand binding sites. Thus, site 42 is located above the substrate access channel in contact with the flexible loop^44-65^ of the B subunit (structural element I) (**Figs. 6** and **S8**). Site 12 is located behind the active site, at the interface of α-helix^230-243^ (named here α1) and α-helix^115-127^ (named here α2) from the A and B subunits, respectively (**Figs. 6** and **7*A***). Last, site 33 is located below the substrate access channel in contact with the α-helix^231-241^ from the B subunit (structural element II) and the α-helix^115-129^ from the A subunit (structural element III) (**Figs. 6** and **8*A***).

**Figure 6.**
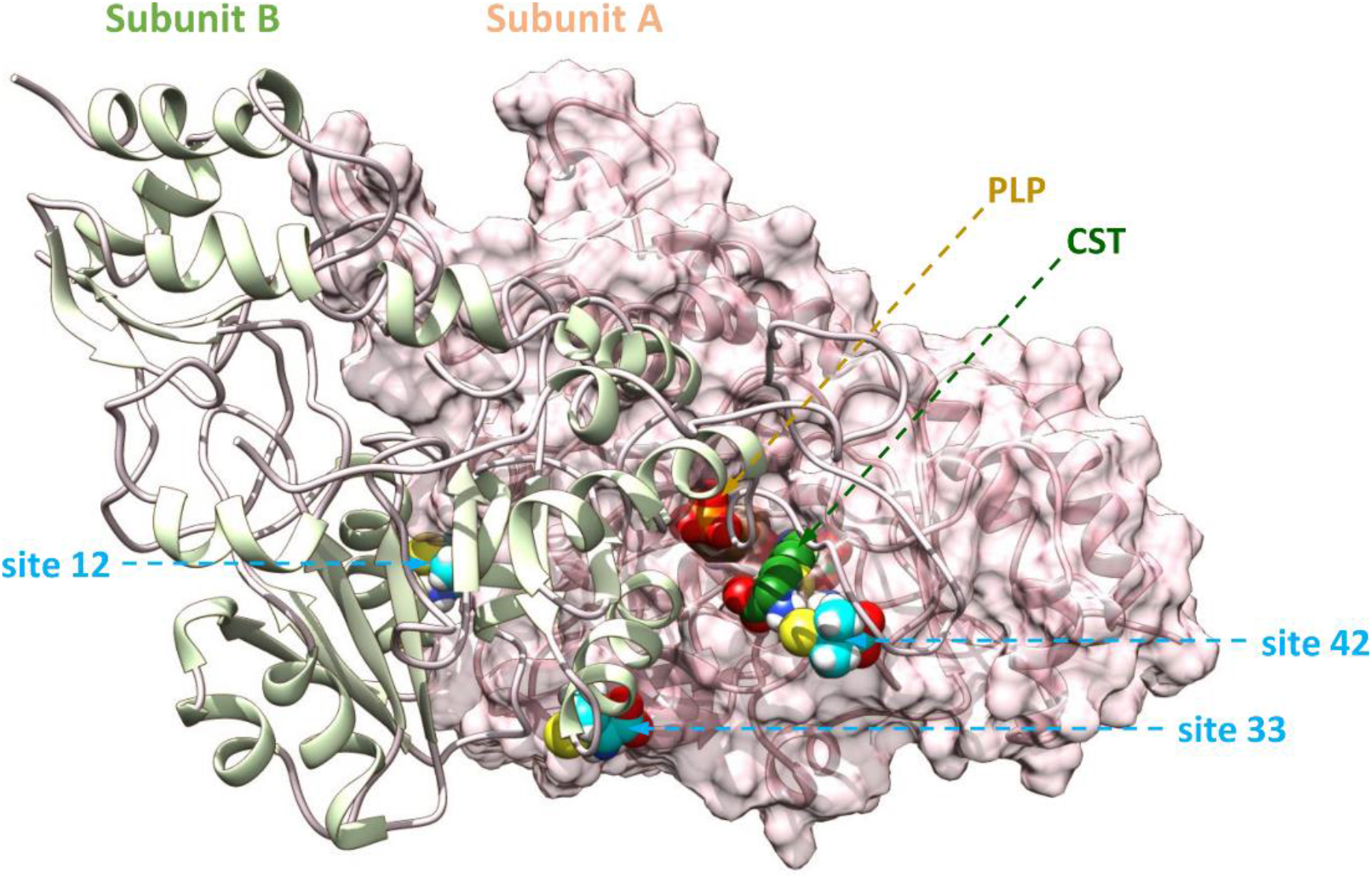
Location of interfacial D-penicillamine binding sites selected for MD simulations. Shown is the position of D-pen (blue sphere) interfacial binding sites. Site 42 is located above the substrate access channel in contact with the flexible loop of the B subunit that contributes amino acid residues to the active site from the A subunit. Site 12 is located behind the active site at more than 20 Å from the PLP cofactor. Site 33 is located below the substrate access channel in contact with α-helices from the A and B subunits which participate in the active site geometry. Cystathionine (CST) and the PLP cofactor are represented in green and brown spheres, respectively.

Thereafter, the binding mode of D-pen was thoroughly examined. Thus, docked D-pen to site 42 (CDOCKER energy and CDOCKER interaction energy of -35.6 kcal/mol and -27.4 kcal/mol, respectively) establishes H-bonds with the backbone of Met354 and the side chains of *Gln49 and Glu339 (**Figs. 6** and **S8**). D-pen also forms C-H…O interactions with the side chain of Met354 and the backbones of *Gly53 and Met354. Docked D-pen to site 12 (CDOCKER energy and CDOCKER interaction energy of -34.1 kcal/mol and -26.7 kcal/mol, respectively) intervenes in salt bridge formation with *Glu127 from α2, in van der Waals contacts with Arg235 and Phe238 from α1, in O…S contact with *Glu127 and in C–H⋯S interaction with Phe238, the later interaction resembling conventional hydrogen bonds (60). Interestingly, while D-pen bound to site 12 is located more than 20 Å away from the active site, the interfacial helix α1 it interacts with establishes hydrogen bond contacts with nearby structural elements (α3 and β1) connected to the active site (**Figs. 6** and **7*A***). Last, docked D-pen to site 33 (CDOCKER energy and CDOCKER interaction energy of -35.7 kcal/mol and -27.4 kcal/mol, respectively) intervenes into H-bond formation with the backbone of Gln123 and is in an H-bond interacting radius with the side chains of Gnl123 and Glu127 (**Fig. S10*A***). In addition, D-pen establishes C-H…O connections with the side chain of *Phe238 and Gln123, van der Waals contact with *Leu239, O…S and C-H…S interactions with Glu127.

**Figure 7.**
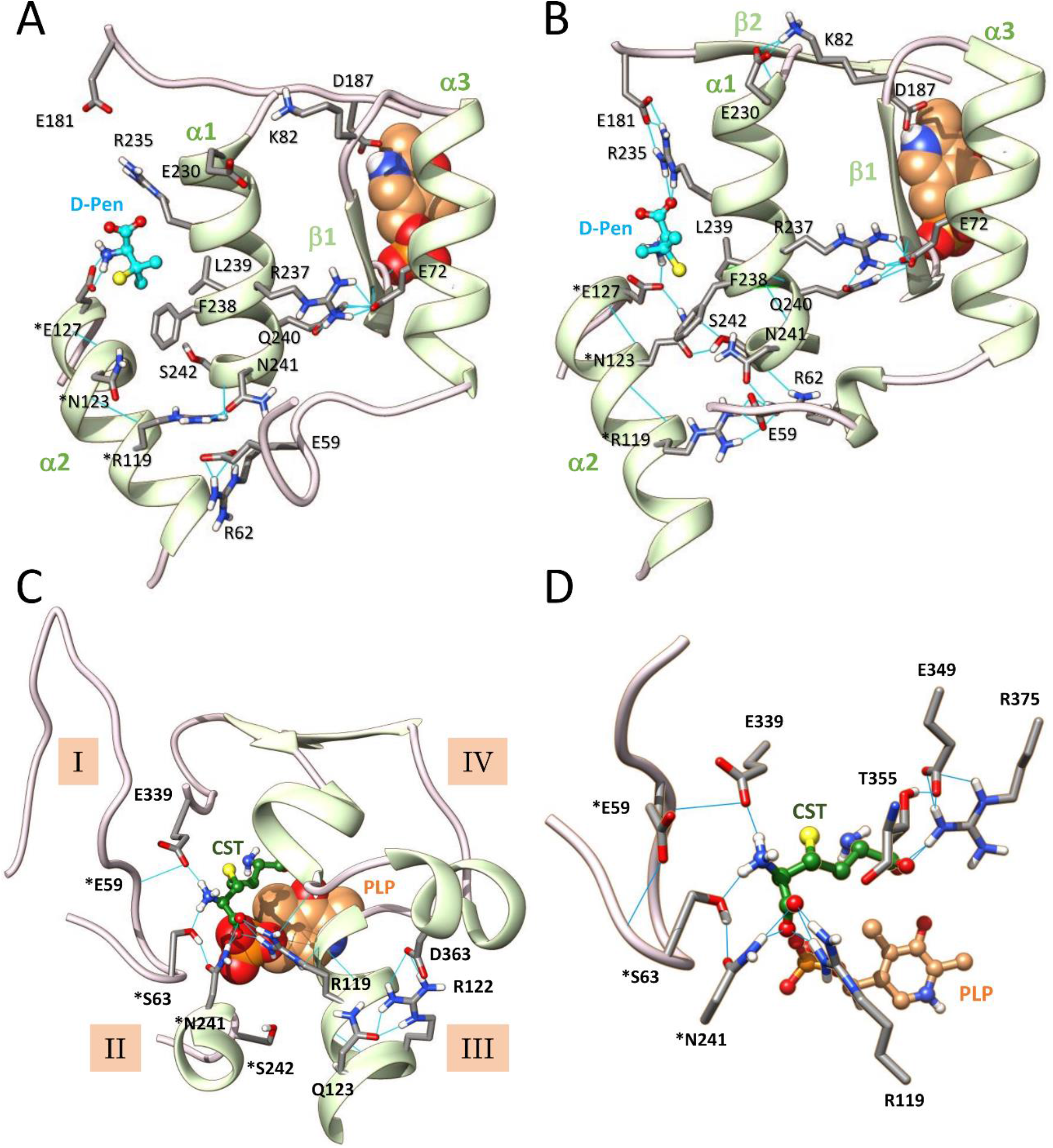
Long range effects of D-penicillamine binding to site 12 propagating to the active site of CSE. (**A, B**) Structural comparison of D-penicillamine (cyan) binding at the interfacial binding site 12 after molecular docking (**A**) or 50 ns MD simulations after (**B**). As shown, MD simulations induce significant changes at the ligand binding site. Thus, a new hydrogen bonding network is created in the vicinity of the interfacial ligand, including new interactions of the interfacial α-helix α1 with its neighbouring. (**C, D**) Representative snapshot of the active site after 50 ns MD simulations showing the hydrogen bonding network partly implicated in the unity of the substrate access channel (**C**) and the effect produced by D-penicillamine binding to site 12 on the cystathionine interacting network (**D**). Particularly, the substrate is retained at the entrance of the substrate access channel by establishing H-bond contacts with *Ser63, *Asn241 and Glu339. Consequently, cystathionine no more establishes an H-bond interaction with the O3’ of PLP and it proximal amino group is now pointing opposite to the C4’ atom of PLP at 4.88 Å away. *Residues from the adjacent subunit B. H-bonding are depicted by light blue lines. CST = cystathionine.

Surprisingly, even though sites 33 and 42 have already been identified in the literature, they have been designated as putative activator binding sites (28, 61). Thus, residues surrounding site 42 have been proposed to establish electrostatic interactions with the phenoxy part of isoxsuprine, a presumed allosteric activator of CSE solely based on high-throughput virtual screening performed with the protein structure in complex with propargylglycine (PDB code: 3COG) (28). However, isoxsuprine extends from the side of site 42 to Thr355 and established electrostatic interaction with Arg119 therefore possibly disturbing the bonding of the substrate to the active site, which raises questions regarding the supposed role of isoxsuprine as an activator of CSE. Importantly, the biochemical validation of a putative isoxsuprine-CSE interaction was not carried out and MD simulations showed that isoxsuprine was not stable during the 10 ns time course of MD simulations, its binding free energy profile strongly fluctuating and becoming positive at the end of the simulations (28). Site 33 on the other hand has been assigned as a possible binding site for the triterpenes ursolic acid and uvaol, designed putative nonpolar allosteric activators of CSE solely based on blind docking studies. Once more, the biochemical validation of the triterpenes-CSE interaction was not performed while the docking score was low (∼3.1 kcal/mol) (61).

### Molecular mechanisms for interfacial inhibition

Next, the best docking pose of each ternary complex CSE-substrate-ligand was subjected to 50 ns MD simulations, not only to investigate their respective stability but also to analyze the structural changes imposed by D-pen binding on the active site and cystathionine positioning. Importantly, the 10-ns all-atom MD simulations of the ternary complex formed with D-pen bound to site 42 show that the protein backbone is rapidly destabilized (**Fig. S4*B***). Although this interfacial ligand binding site is quite interesting due to its location nearby the active site and even if we cannot entirely exclude it as a likely ligand binding site, we did not consider this complex afterwards. Both the substrate and the ligand bound to sites 12 and 33 are stable during the molecular dynamics, but the ternary complexes lose the initial stability provided by substrate binding as suggested by an increase in RMSD (ΔRMSD = 0.21 Å and 2.14 Å over the last 20 ns for site 12 and site 33 *vs* the binary complex, respectively) and RMSF (ΔRMSF = 0.30 Å and 0.81 Å for site 12 and site 33 *vs* the binary complex, respectively) in the ternary complexes compared to the binary complex (**Fig. S4*C-E***).

Interestingly, MD simulations induce changes in the immediate vicinity of the interfacial D-penicillamine binding sites both at sites 12 (**Fig. 7*B***) and 33 (**Fig. 8*A***). In site 12, D-pen now becomes stabilized through H-bonding with *Glu127 and the NH2 and Nε of Arg235, van der Waals contact with Arg235, unconventional O…S and C-H…S interactions with *Glu127 and Phe238/Leu239, respectively. Interestingly, new connexions are established between α1 and β2 which carries Asp187 itself H-bonded to the pyridine ring of PLP, and between α1 and the α3-β1 part which ends with amino acid residues interacting with the phosphate group of the cofactor. These results suggest that structural changes provoked at the interfacial ligand binding site could translate to the active site through these new connexions (**Fig. 7*B***). In site 33, D-pen establishes H-bonds with Glu127 and *Arg235, C-H…O contact with Glu127 and its methyl groups participate in van der Waals interactions with *Arg235, *Phe238 and *Leu239 (**Figure 8*A***). Importantly, many perturbations recorded in the CSE-substrate-ligand ternary complexes during the 50 ns MD simulations are not observed at both site location in the CSE-substrate binary complex, thus revealing that binding of D-pen at either site 12 or 33 induces the creation of an entirely new interacting network at the vicinity of the interface between the A and B subunits (**Figs. S9** and **S10**). Interestingly, interface alteration has also been observed in the context of benzothiazole derivative inhibitors of triosephosphate isomerase (TIM) from *Trypanosoma crizi* (62). Thus, these interfacial inhibitors produce an allosteric effect on TIM catalytic activity by producing a constrained region at subunit interface through enhancement of intersubunit H-bonds, changing the solvent exposure of catalytic residues and altering the collective dynamics of the enzyme (62).

**Figure 8.**
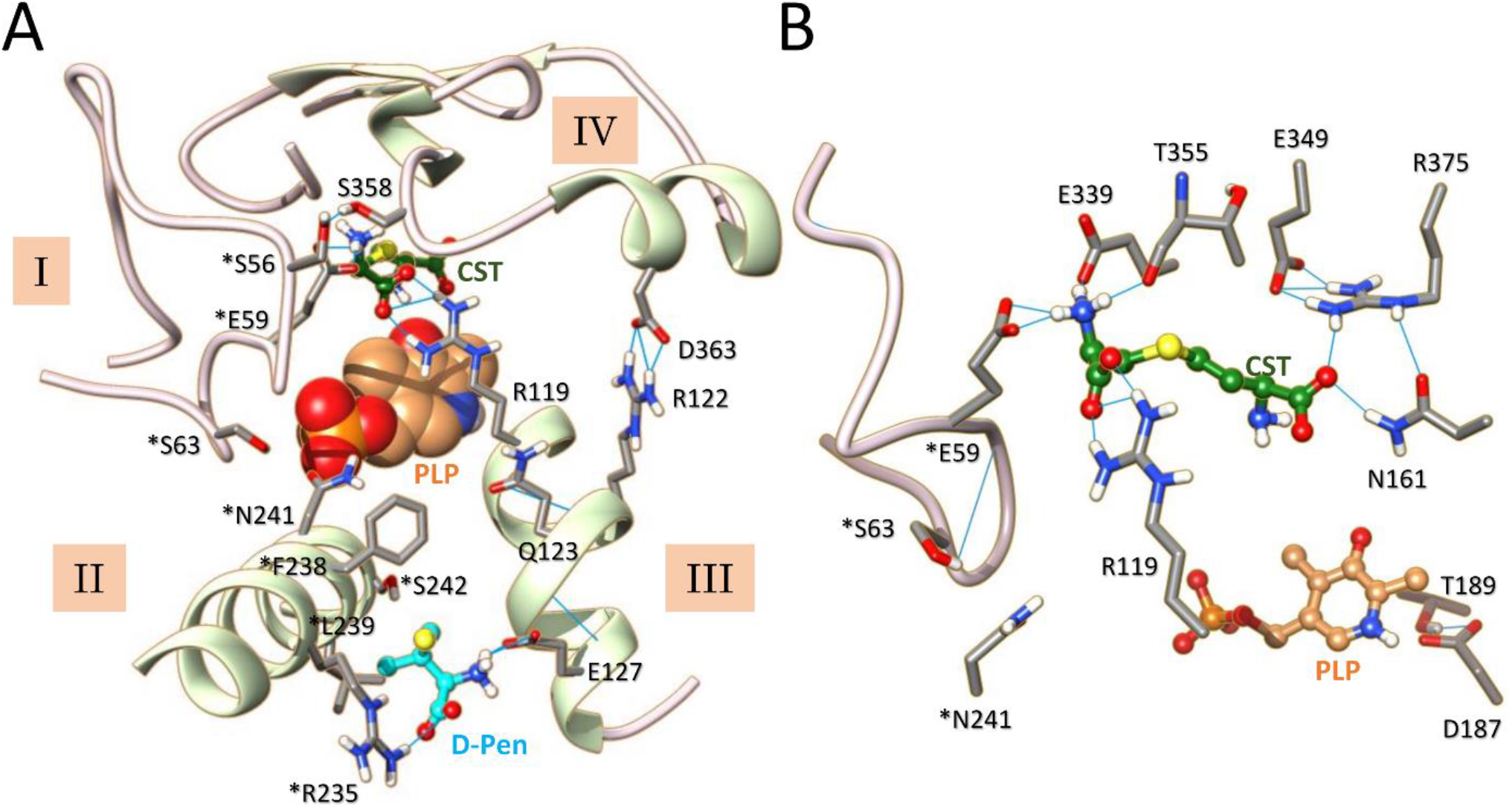
Destabilization of CSE active site by D-penicillamine binding at site 33 during molecular dynamics (MD). (**A, B**) Representative snapshot of the active site after 50 ns MD simulations showing the impact of D-pen (cyan) binding on the hydrogen bonding network partly implicated in the unity of the substrate access channel (**A**) and the cystathionine interacting network (**B**). Significantly, binding of D-pen to the interfacial ligand-binding site located between the α-helices from structural elements II and III causes significant disturbances in the opening of the substrate access channel (**A**) and important variations at the active site, such as the stabilization of the distal ammonium group of cystathionine by H-bonding with *Glu59 and Glu339 and the disruption of the H-bond contact between the pyridine ring of the cofactor and Asp187 (**B**). *Residues from the adjacent subunit. H-bonding are depicted by light blue lines. CST = cystathionine.

MD simulations also provide evidence for long range effects of inhibitor binding propagating to the active site with the initial hydrogen bonding network connecting structural elements I-IV being profoundly perturbed (**Figs. 7*C*** and **8*A***), as are the cofactor (**Fig. S6**) and substrate (**Fig. S7**) interacting networks. Significantly, structural elements II and III are no more directly connected while structural elements I and IV are now connected through H-bonding between *Glu59 and Glu339 (site 12) (**Fig. 7*C***) or between *Ser56 and Ser358 (site 33) (**Fig. 8*A***). This results among other things in the opening of the substrate access channel in comparison to the binary complex (**Table S1**). At the active site, the PLP cofactor is destabilized either at its phosphate group (site 12) or at its pyridine moiety (site 33) thus explaining our fluorescence spectroscopy results which show that the attachment of D-pen to the CSE disturbs the intrinsic fluorescence of the PLP probe (**Fig. 2**). Thus, the phosphate group of the cofactor no longer establishes hydrogen bonds with the polypeptide chains of Gly90 and Leu91 when D-pen is bound to site 12, and the pyridine ring no longer forms H-bond with Asp187 when D-pen is bound to site 33. Consequently, the PLP can no more be activated by N-protonation in the latter situation (63). Accordingly, the PLP moiety loses its net positive charge that has been estimated mandatory to initiate catalysis (64). Besides, Tyr114 falls within a radius of hydrogen interaction with O2P and O4P atoms from the PLP cofactor (**Fig. S6**). In addition, the microenvironment in which the tyrosine hydroxyl group was originally found (interactions with the S atom of the substrate) no longer exists therefore rendering Tyr114 unable to participate in a putative proton transfer reaction towards the leaving group of the substrate (**Fig. S7**). Last, in the presence of D-pen, the substrate does not anymore slide deeper into the substrate access channel due to direct H-bonding with amino acid residues located at the entrance of the substrate access channel such as *Ser63, *Asn241 and Glu339 (site 12) (**Fig. 7*D***) or *Glu59 and Glu339 (site 33) (**Fig. 8B**). As a result, the connections initially established between cystathionine and the surrounding amino acids are completely disrupted and the distances between its nucleophilic amino group and the O3’ or C4’ atoms of the PLP increases significantly (**Fig. S7** and **Table S1**), no longer leaving the possibility for the nucleophilic group of the substrate either to establish an H-bond with the O3’ atom or to attack the C4’ position of the PLP in order to form the *gem* diamine intermediate. Altogether, our observations indicate that sites 12 and 33 are interesting target sites for the design of selective allosteric inhibitors that will become potent pharmacological modulators of CSE functions

## Conclusion

Our findings provide a comprehensive picture of the structure and dynamics of cystathionine binding to CSE, and how they are modulated by D-penicillamine binding. Our modeling studies thus reveal that cystathionine binding promotes the organizing of the active site prior to catalysis, in particular by confining the access channel from the substrate to the active site and by establishing various interactions with surrounding amino acid residues including S/π and O…S interactions with between its S atom and Tyr114 that could therefore intervene in protonation of the leaving group. Strikingly, the proximal amino group of cystathionine also forms a H-bond contact with the O3’ atom of the PLP cofactor, thus suggesting that the O3’ atom could participate in proton transfer reactions and/or increase the selectivity of the nucleophilic substrate amino group and modulate the electrophilic character of the C4’ atom. Interestingly, our biochemical analyses reveal that D-penicillamine acts as a peculiar inhibitor of both cystathionine cleavage and H_2_S biogenesis by human CSE by adopting an unusual mixed inhibition mechanism. Our MD simulations disclose that D-penicillamine binds to the interface of two adjacent subunits with inhibitor binding inducing the making of an entirely new interacting network at the vicinity of the interface between enzyme subunits. Remarkably, this interface modulation propagates to the active site and promotes an inactive state of the enzyme by perturbing PLP-enzyme, substrate-enzyme, and PLP-substrate interactions, thus explaining D-pen mode of inhibition. Overall, our studies should not only help for the design of new allosteric interfacial inhibitors against H_2_S biogenesis by cystathionine γ-lyase but also facilitate the targeted virtual high-throughput screening of libraries of putative allosteric compounds.

## ^1^Abbreviations

TSP: transsulfuration pathway
PLP: pyridoxal 5’-phosphate
CBS: cystathionine β-synthase
CSE: cystathionine γ-lyase
CST: cystathionine
D-pen: D-penicillamine
PDB: protein data bank
RMSD: root mean square deviation
RMSF: root mean square fluctuation

## Author contributions

LLC performed computational studies and edited the manuscript; DP realized biochemical experiments, analyzed data, and wrote the manuscript.

## Acknowledgments

D.P. thanks Dr E. Galardon (Université Paris Cité, CNRS, UMR 8601) for useful discussions and editing of the manuscript. The authors would like to acknowledge the Molecular Modeling Platform of Université Paris Cité⎮CNRS⎮UMR 8601 for the computational studies.

## Supplementary Materials

The following supporting information are available, Table S1: Effect of D-pen on d(substrate-O3’) or d(substrate-C4’) and on the perimeter of the substrate access channel entrance; Scheme S1: Mechanism and characteristics of mixed inhibition; Scheme S2: Implication of the O3’ atom from PLP into proton transfer; Figure S1: Purification of recombinant human cystathionine γ-lyase; Figure S2: Influence of the N-terminal 6xHis-tag on CSE activity; Figure S3: Inhibition of CSE-catalyzed H_2_S or cysteine production by D-penicillamine; Figure S4: RMSD and RMSF plots; Figures S5: Interacting network of PLP and cystathionine from molecular docking; Figures S6-S7: Structural comparison of the impact of D-pen binding to CSE on the interacting network of PLP or cystathionine after 50 ns MD simulations; Figure S8: Representative docking pose for the binding of D-pen at site 42; Figures S9-S10: Structural comparison of D-penicillamine binding at site 12 or 33 after docking and MD simulations.

## Supporting Information

**Table S1.**
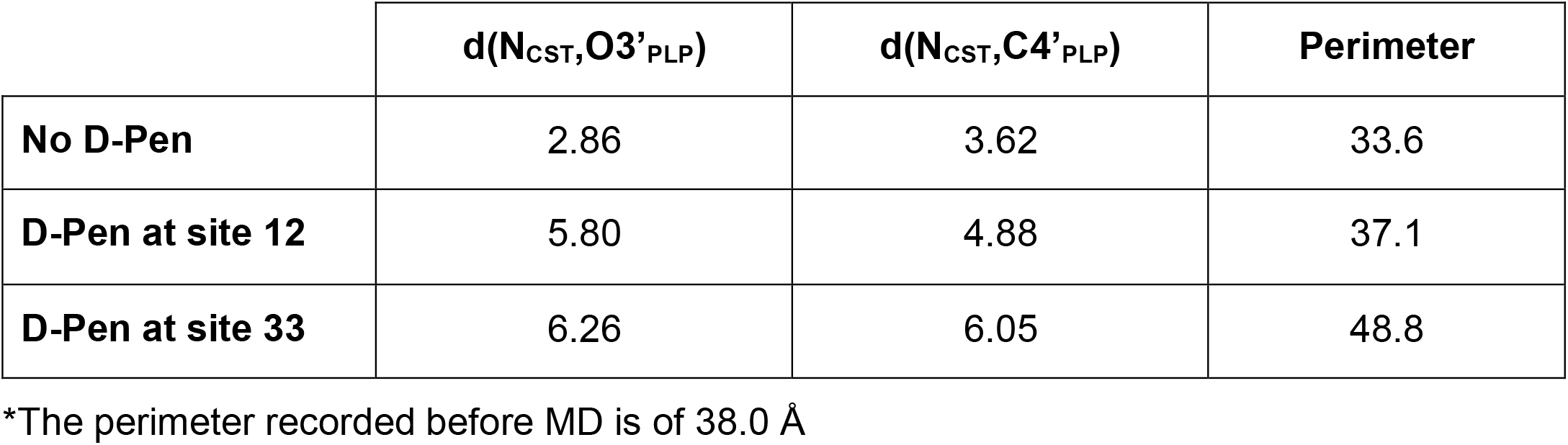
Effect of D-pen on the distances d(N_CST_,O3’_PLP_), d(N_CST_,C4’_PLP_) and on the perimeter* of the substrate access channel entrance (in Å) recorded after 50 ns MD simulations.

|  | **$d(N_{CST}, O3'_{PLP})$** | **$d(N_{CST}, C4'_{PLP})$** | **Perimeter** |
| --- | --- | --- | --- |
| **No D-Pen** | 2.86 | 3.62 | 33.6 |
| **D-Pen at site 12** | 5.80 | 4.88 | 37.1 |
| **D-Pen at site 33** | 6.26 | 6.05 | 48.8 |

\*The perimeter recorded before MD is of 38.0 Å

**Scheme S1.**
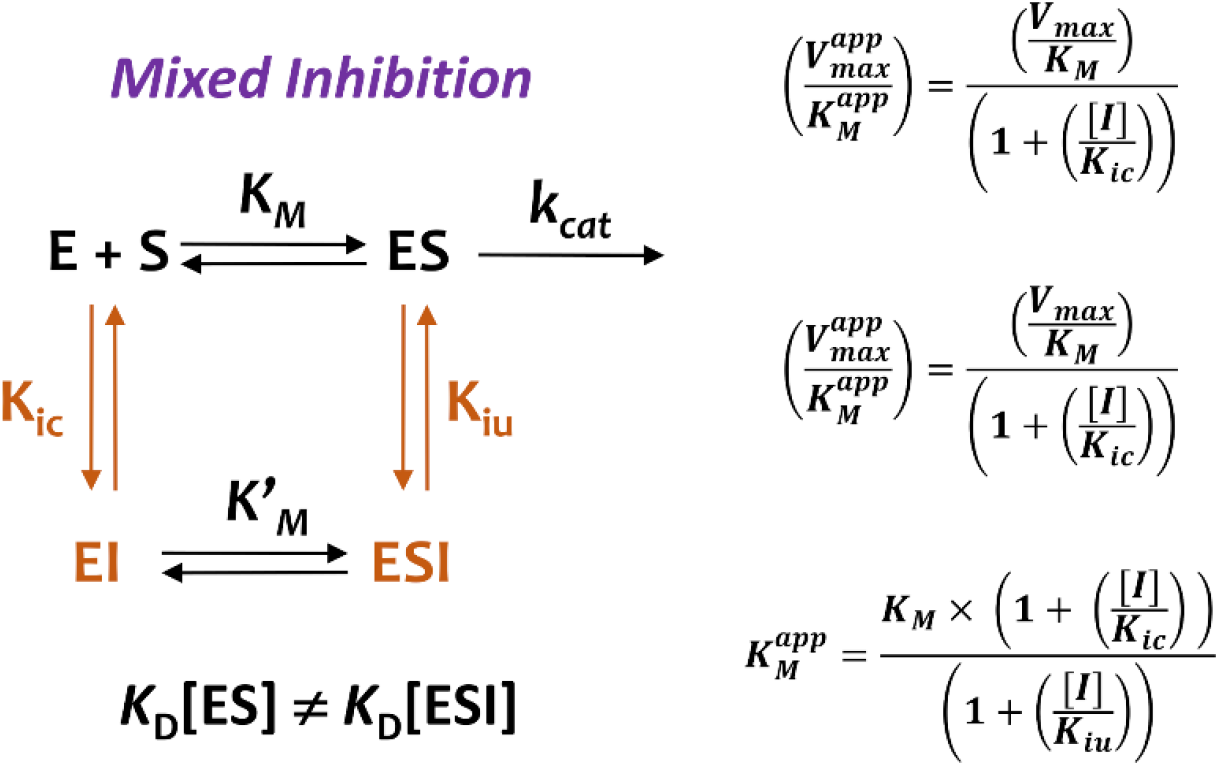
Mechanism and characteristics of mixed inhibition. E, enzyme; S, substrate; ES, Michaelis complex; I, inhibitor.

**Scheme S2.**
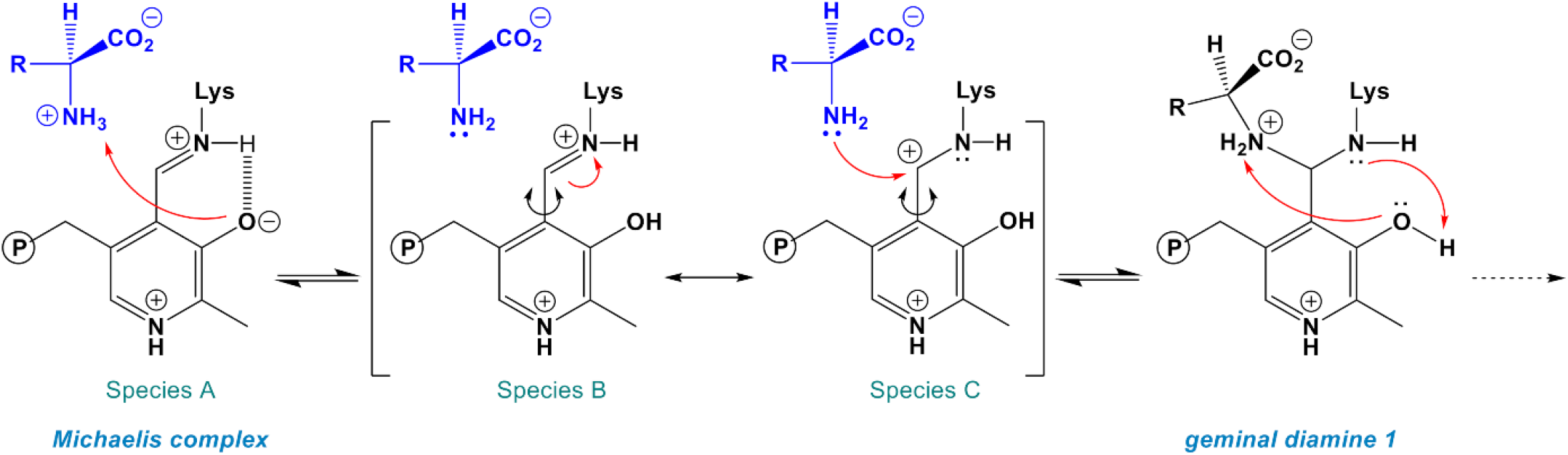
Implication of the O3’ atom from PLP into proton transfer. Catalytic role of O3’ atom in the proton transfer from the substrate (blue) to the lysine amino group from the Schiff base linkage, as postulated for the L-serine hydratase from *Xanthomonas oryzae* pv. *Oryzae*.

**Figure S1.**
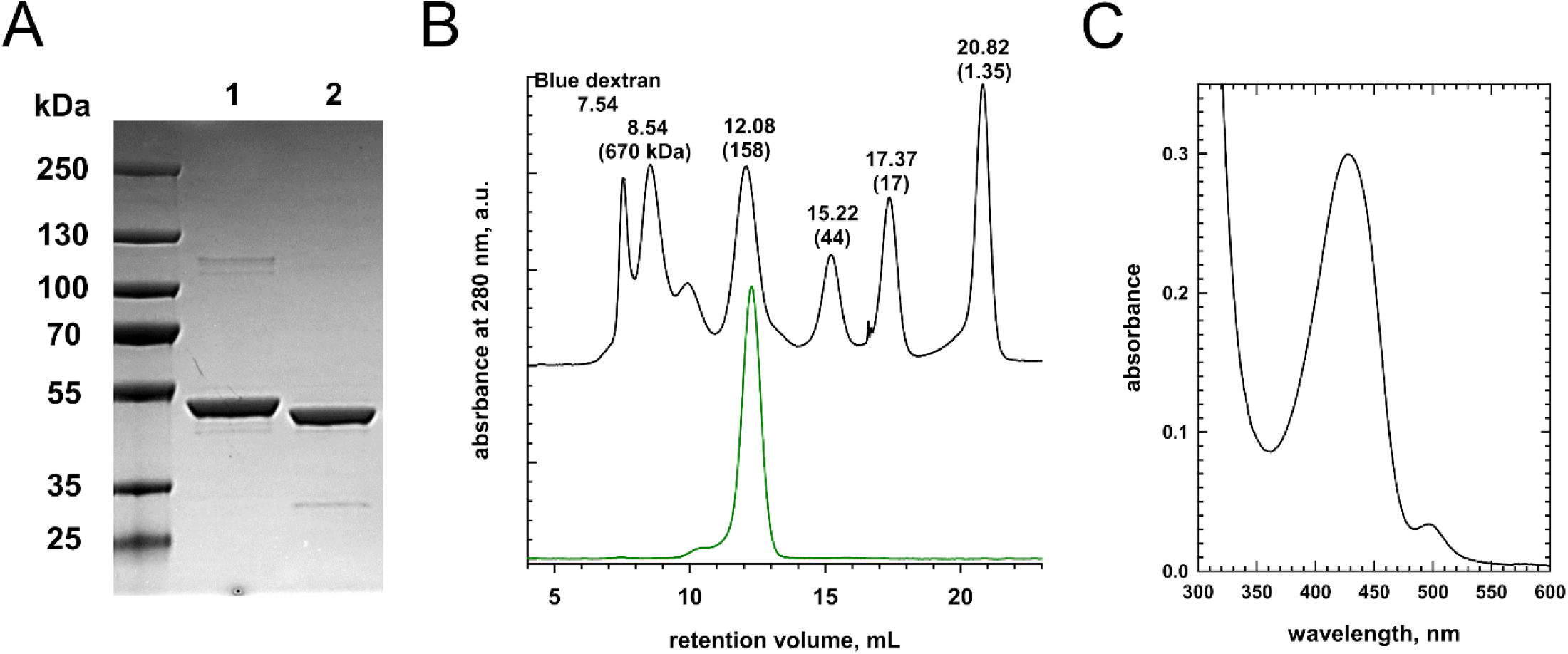
Purification of recombinant human cystathionine γ-lyase. (**A**) 12% SDS-polyacrylamide gel of recombinant human CSE (12 μg) before (**1**) and after (**2**) cleavage of its N_terminal_ 6xHis-tag by TEV protease. The molecular masses of the Precision Plus Protein standards (left lane, Bio-Rad) are shown. (**B**) Gel filtration analysis of purified recombinant tag-less human CSE (100 μg - green line) on a 10/300 GL Superdex-200 size exclusion column (GE Healthcare) eluted with 100 mM HEPES, 150 mM NaCl at pH 7.4 at 0.5 mL/min. The elution profile of (Bio-Rad) protein standards (black line) separated under the same conditions is shown for comparison. The apparent retention volume and molecular weights of protein standards used are labelled on the plot. The latter are thyroglobulin (670 kDa), bovine γ-globulin (158 kDa), chicken ovalbumin (44 kDa), equine myoglobin (17 kDa), and vitamin B_12_ (1.35 kDa). The retention volume of tag-less human CSE (12.220 ± 0.062 mL, n=2±SD) corresponds to an apparent molecular weight of 148 ± 4 kDa. Under similar conditions, full-length CSE eluted with a retention volume of 12.157 ± 0.045 mL (n=2±SD), which corresponds to an apparent molecular weight of 152 ± 3 kDa. (**C**) UV-Visible spectrum of the PLP cofactor of purified recombinant human cystathionine γ-lyase monitored in 100 mM HEPES at pH 7.4 and 20°C. The internal aldimine form of the PLP cofactor exhibits a sharp band at 428 nm. A small proportion of purified CSE contains PLP-aminoacrylate species that displays an absorption peak around 495 nm.

**Figure S2.**
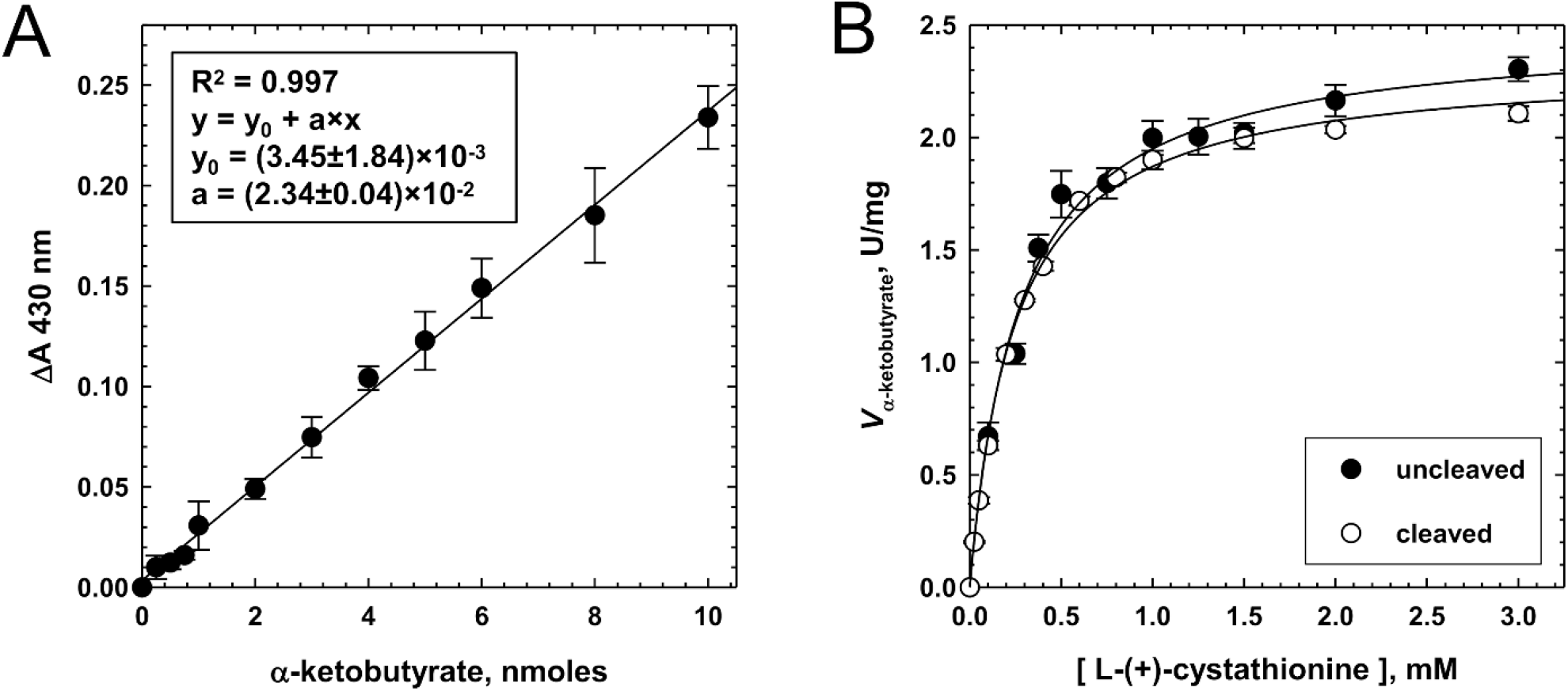
Influence of the N-terminal 6xHis-tag on CSE activity. (**A**) Etalon curve for the determination of α-ketobutyrate content in the 2,4-dinitrophenyl hydrazine test (n = 4 ± SD). (**B**) Kinetics of α-ketobutyrate generation from cystathionine by CSE (uncleaved) and 6xHis-tag-less CSE (cleaved) (n = 2 ± SD). Cysteine synthesis from L-(+)-cystathionine was determined by measuring α-ketobutyrate production at 37°C from assay mixtures containing 0.1-4 mM L-(+)-cystathionine and CSE (10 μg) in 100 mM HEPES at pH 7.4. Hyperbolic fit to the data gave V_max_ and *K*_M_ values of 2.47 ± 0.05 U/mg and 0.269 ± 0.020 mM for CSE, and 2.33 ± 0.02 U/mg and 0.246 ± 0.006 mM for tag-less CSE.

**Figure S3.**
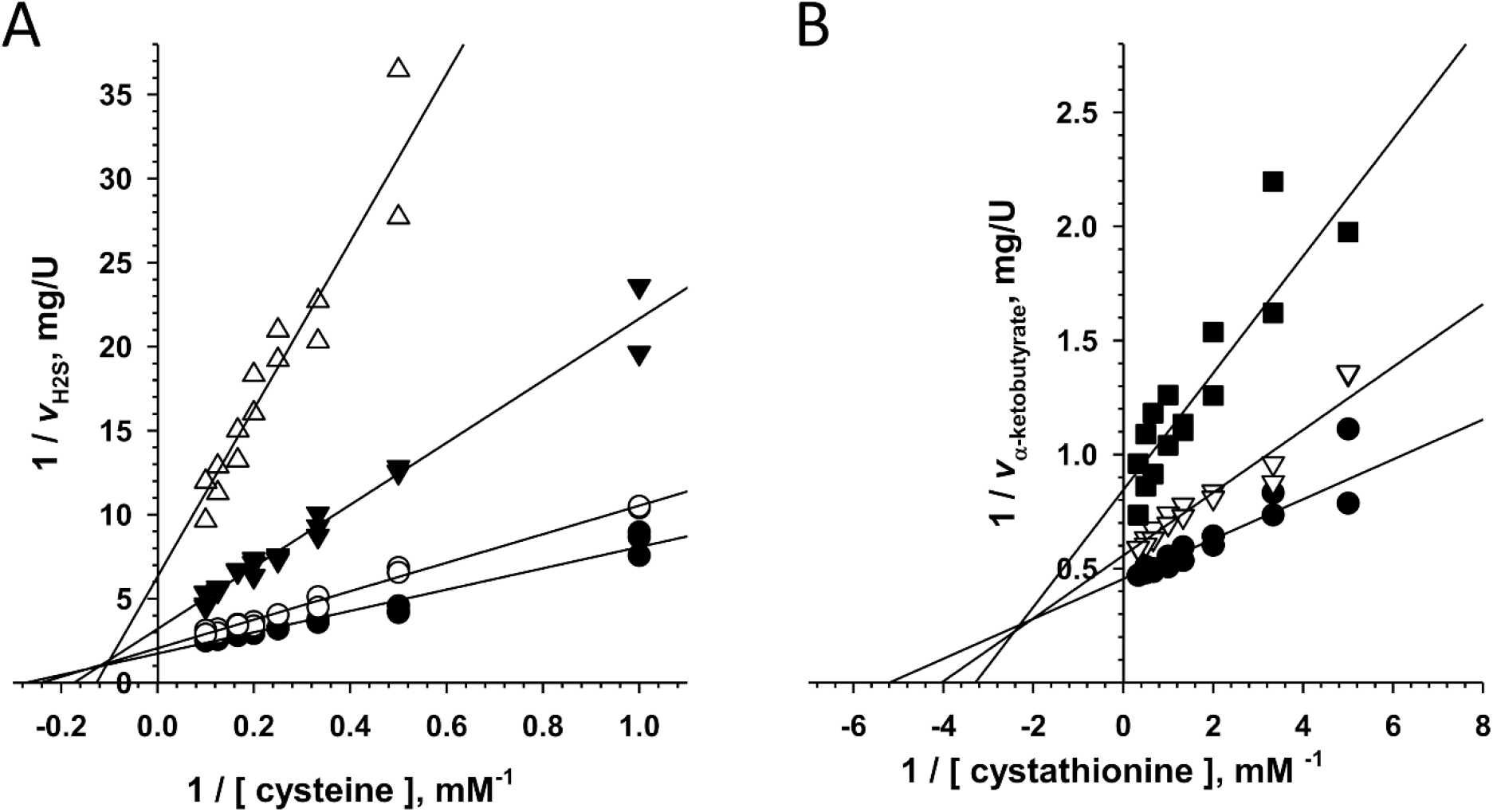
Inhibition of H_2_S production and cystathionine cleavage by D-penicillamine. (**A**) Lineweaver-Burk plot analysis of CSE inhibition by D-penicillamine (n ≥ 2 ± SD) as determined in **Fig.3B**. Assay mixtures contained 1-10 mM cysteine, 0.4 mM lead acetate, CSE (40 μg) and varying concentrations of D-pen (⬤, 0 mM; ○, 1 mM; ▼, 5 mM; △, 10 mM) in 100 mM HEPES at pH 7.4 and 37 °C. (**B**) Lineweaver-Burk plot analysis of CSE inhibition by D-penicillamine (n = 2 ± SD) as determined in **Fig.4A**. Assay mixtures contained 0.1-3 mM cystathionine, CSE (5 μg) and varying concentrations of D-pen (⬤, 0 mM; ▽, 2 mM; ⬛, 5 mM) in 100 mM HEPES at pH 7.4 and 37 °C.

**Figure S4.**
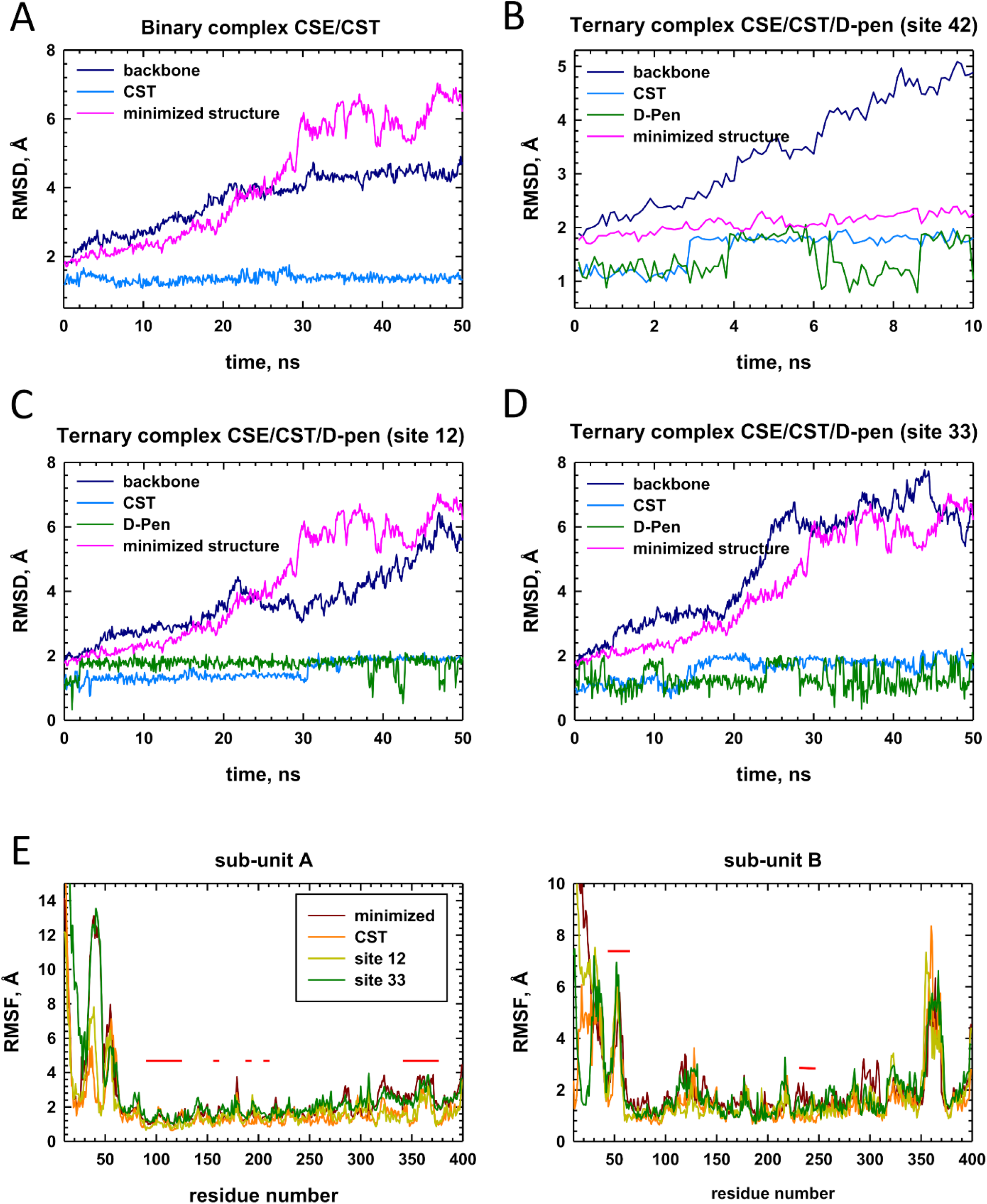
RMSD and RMSF plots. (**A-D**) Root mean square deviation (RMSD) analysis was carried out for the molecular dynamics simulations of each system (top and middle plots). The RMSD analysis of the minimized structure is shown in each plot for comparison. (**E, F**) The RMSF (for the backbone of each residue) of minimized, binary and ternary complexes trajectories were calculated along the entire sampling interval (50 ns). The horizontal red lines indicate the position of amino acids that are integral to the active site and participate in the substrate channel unity. An analysis of the RMSD of CST and D-pen show that substrate and ligand are both stable in their respective binding pocket along the entire dynamics range (50 ns). Importantly, only active site residues belonging to the flexible loop^44-65^ from the B subunit (structural element I - see Fig. 5*A*) seem to display some flexibility during molecular dynamics simulations. Reduced RMSD value of the enzyme and reduced RMSF of the enzyme residues observed for the binary CSE-CST complex indicate that substrate binding stabilizes the complex.

**Figure S5.**
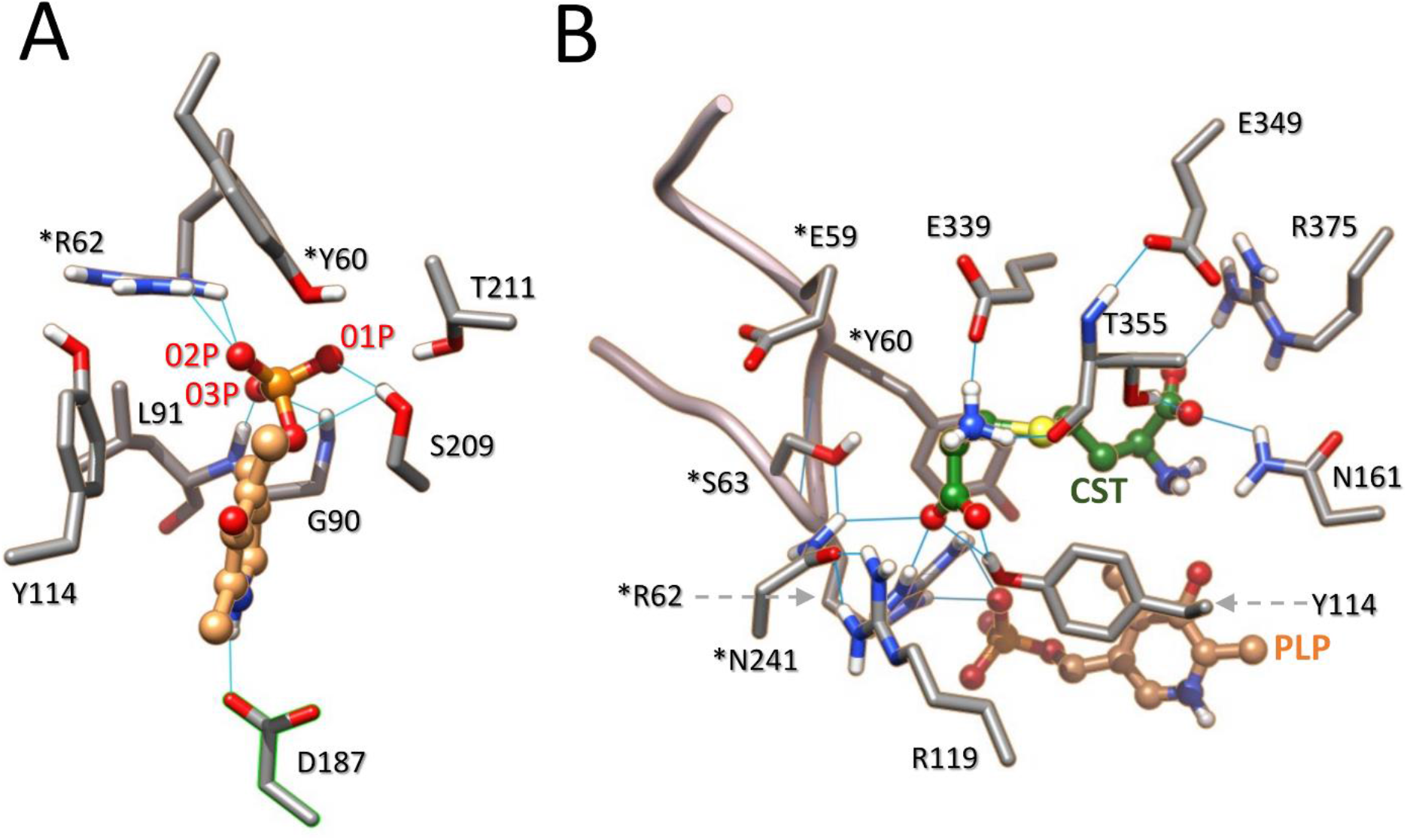
Interacting networks of PLP and cystathionine from molecular docking. Representative docking poses showing the interacting network of PLP (**A**) and cystathionine (**B**) in the active site from the A subunit. The imine bond between the C4’ atom of PLP and Lys212 has been omitted for clarity. *Residue from the adjacent B subunit. H-bonds are depicted by light blue lines.

**Figure S6.**
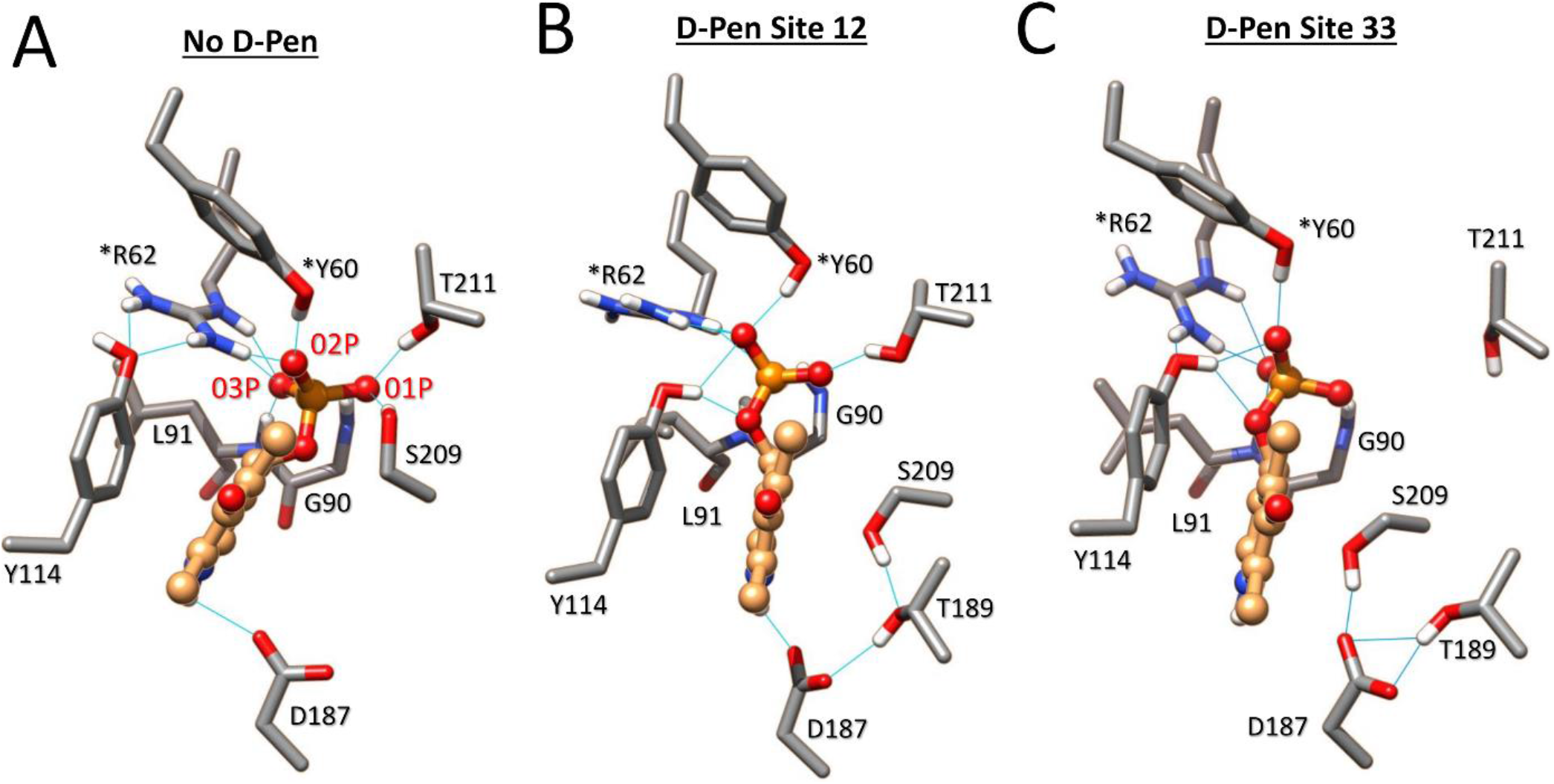
Structural comparison of the impact of D-pen binding to CSE on the interacting network of PLP after 50 ns MD simulations. Structural comparison of the H-bond network of PLP in the active site from the A subunit during MD at t=50 ns in the absence (**A**) or the presence of D-penicillamine bound to site 12 (**B**) or 33 (**C**). Significantly, the PLP cofactor is destabilized at its phosphate group and/or its pyridine moiety when D-penicillamine is bound to site 12 or site 33. Thus, the phosphate group of the cofactor no longer establishes H bond contacts with the backbone of Gly90 and Leu91 when D-pen is bound to site 12 while the PLP is no more stabilized by H-bonding with Thr211 and no more activated by N-protonation in the presence of D-Pen at site 33. In addition, Tyr114 falls within a radius of hydrogen interaction with O2P and O4P atoms from PLP in the presence of the ligand. *Residues from the adjacent subunit B. H-bonding are depicted by light blue lines.

**Figure S7.**
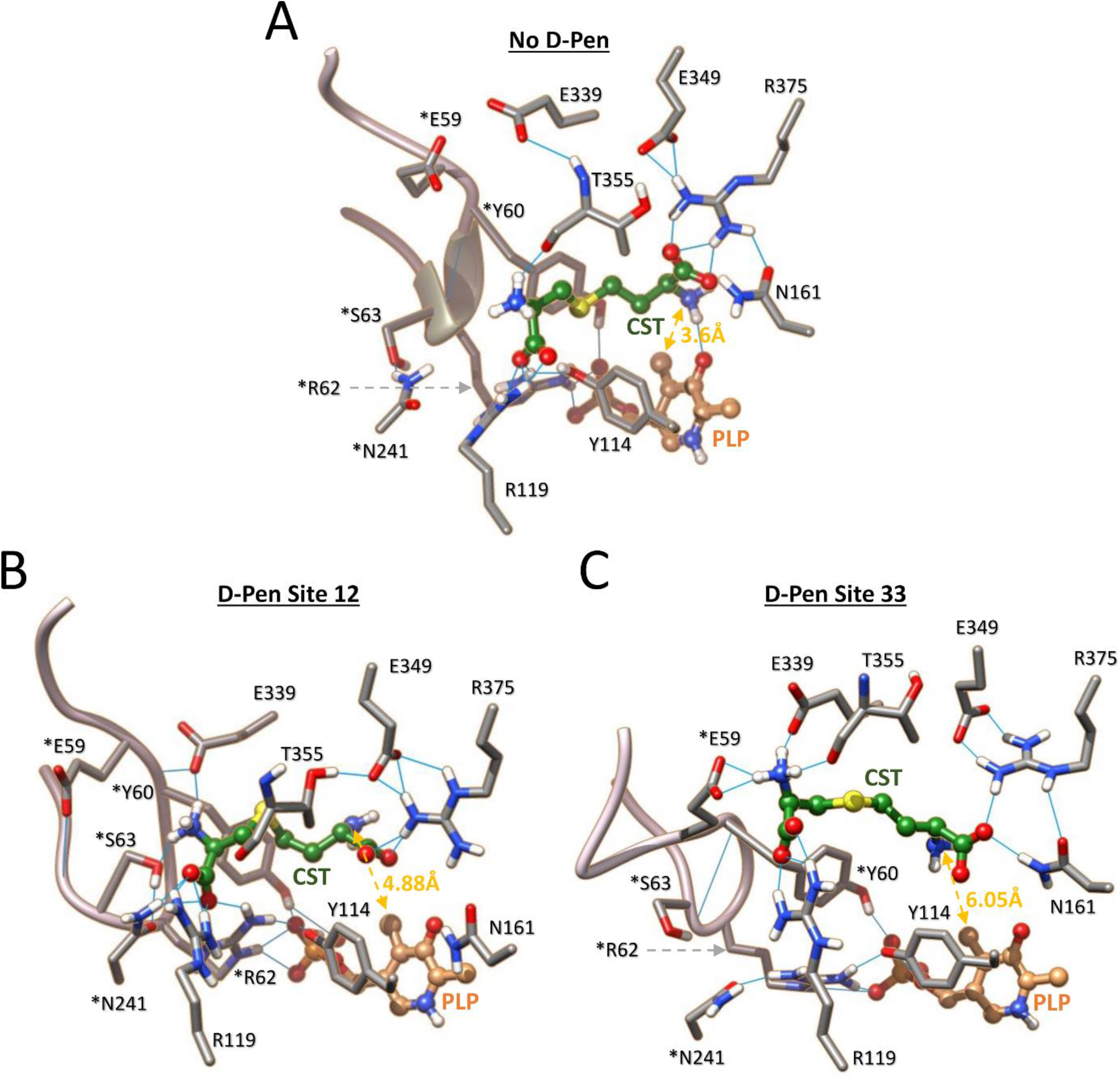
Structural comparison of the impact of D-pen binding to CSE on the interacting network of cystathionine after 50 ns MD simulations. Structural comparison of the interacting network of cystathionine in the active site from the A subunit in the absence (**A**) or the presence of D-penicillamine bound to site 12 (**B**) or 33 (**C**). Importantly, the substrate establishes H bond contacts with amino acid residues located at the entrance of the substrate access channel in the presence of D-pen, *i*.*e*. Glu339, *Ser63 and *Asn241 (site 12) or Glu339 and *Glu59 (site 33). In addition, the interacting network of cystathionine is disrupted with the distances (in yellow) between its nucleophilic amino group and the O3’ or C4’ atoms of the PLP significantly increased. *Residues from the adjacent subunit B. H-bonding are depicted by light blue lines.

**Figure S8.**
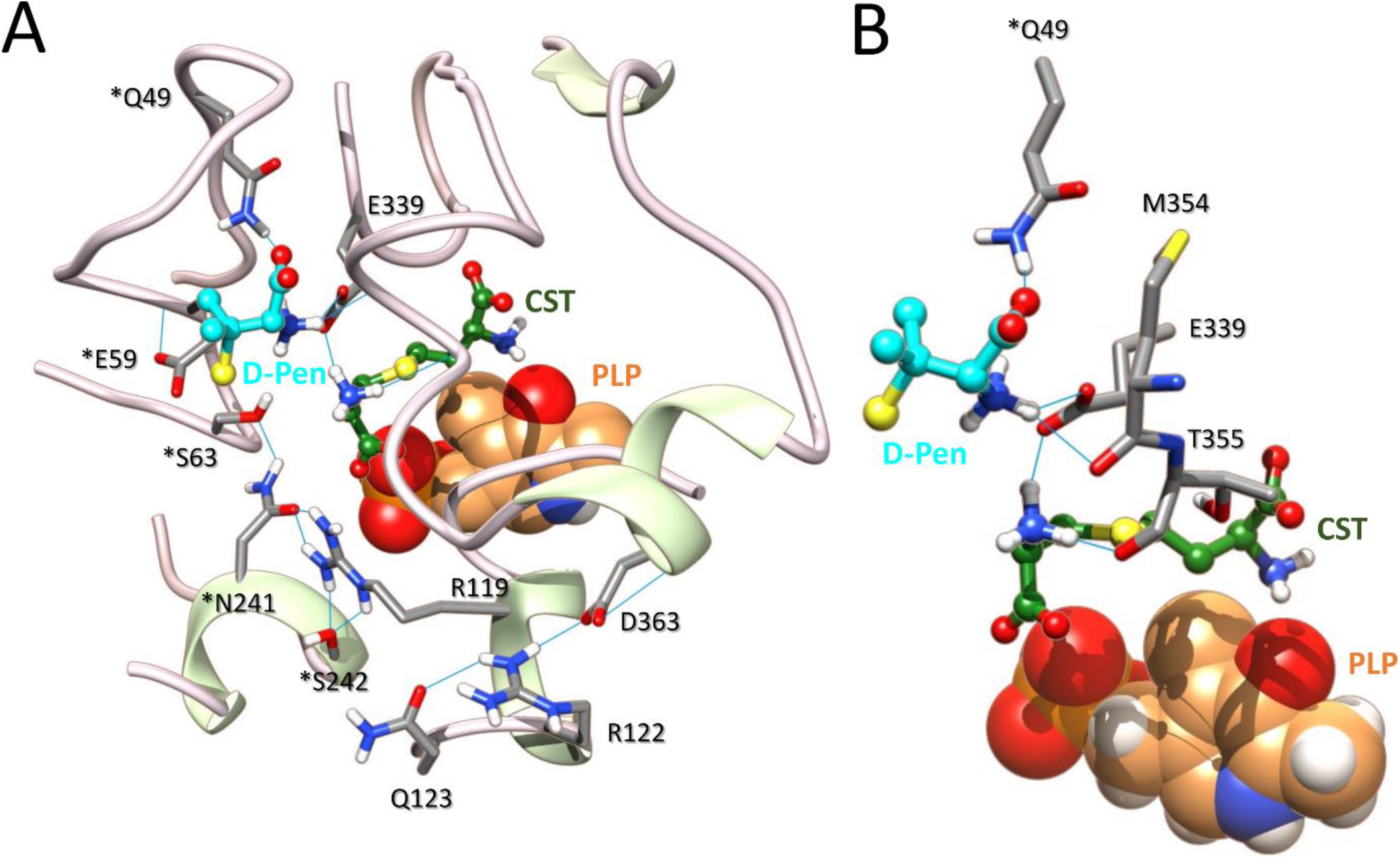
Representative docking pose for the binding of D-penicillamine at the interfacial binding site 42. (**A, B**) Structure of a representative docking pose showing the binding location of D-pen at site 42 (**A**) and close-up of the same docking pose showing the interacting network of the ligand (**B**). As shown, D-penicillamine binds on top of the entrance of the substrate access channel from the A subunit and establishes amongst others H bond contact with Glu339 which is also implicated in substrate stabilization. *Residues from the adjacent subunit B. H-bonding are depicted by light blue lines.

**Figure S9.**
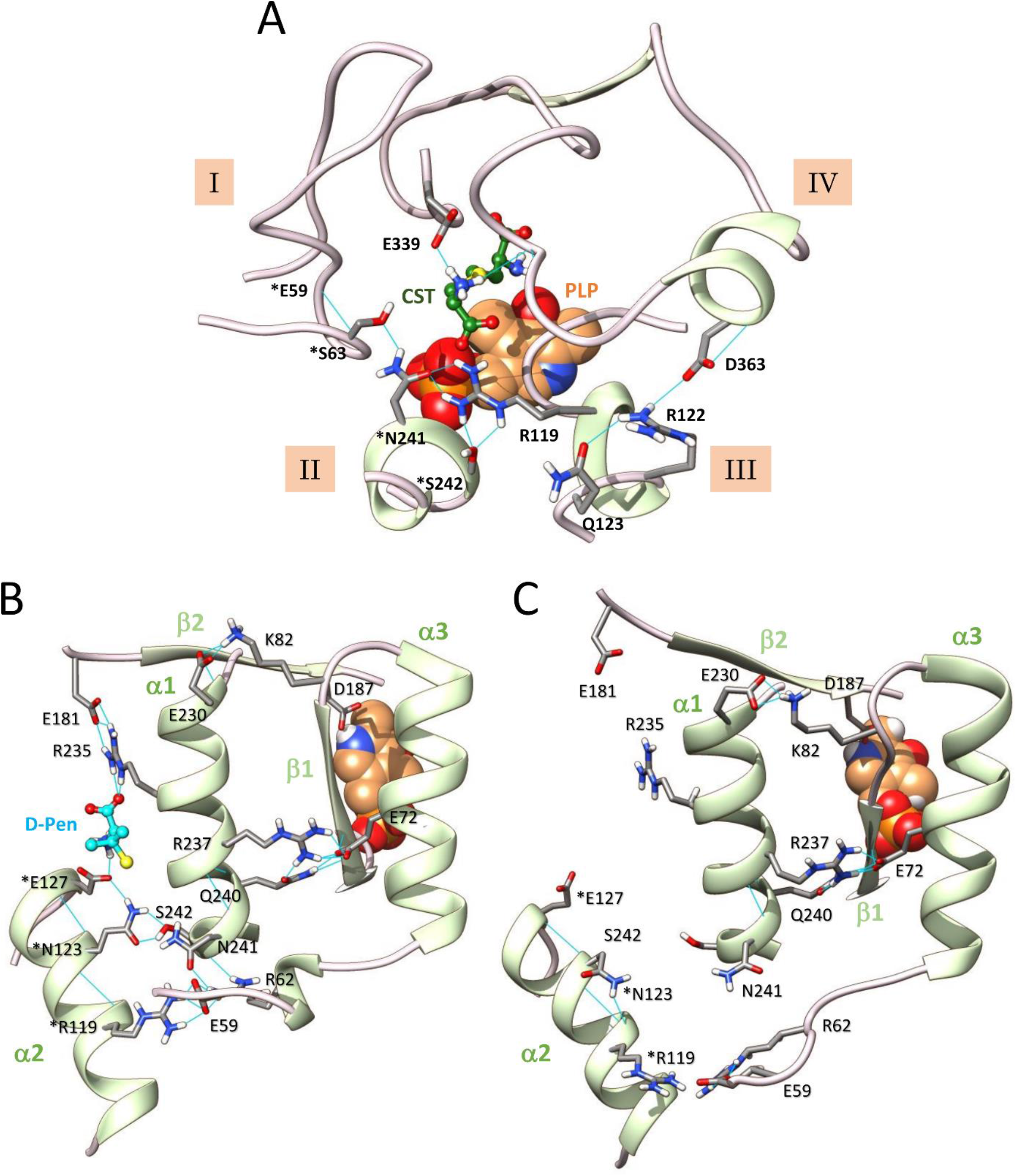
Structural comparison of D-penicillamine binding at site 12 after docking and MD simulations. (**A**) Representative docking pose of the active site when D-pen is bound to site 12. (**B, C**) Structural comparison of representative snapshots obtained after 50 ns of molecular dynamics of the interfacial D-pen binding site 12 in the ternary (**B**) or binary (**C**) complexes. As shown, the comparison of both structures reveal that D-pen binding between interfacial α-helices α1 and α2 promotes substantial structural changes in the vicinity of the interface. * Residue from the adjacent B subunit B. H-bonds are depicted by light blue lines.

**Figure S10.**
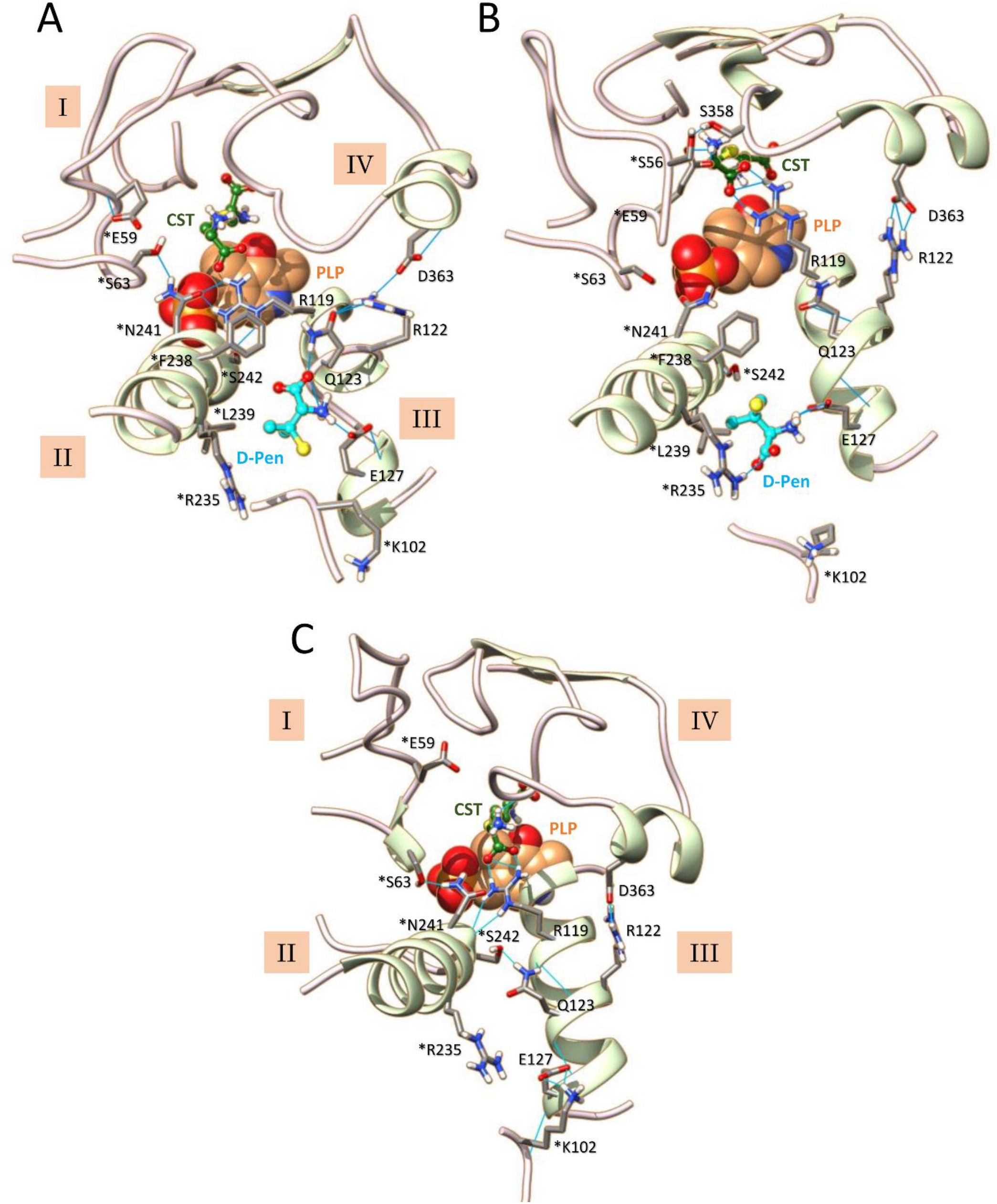
Structural comparison of D-penicillamine at site 33 after docking and MD simulations. (**A**-**C**) Structural comparison of the active site from the A subunit in the presence of D-pen at site 33 after molecular docking (**A**) and 50 ns MD simulations of the ternary CSE-substrate-ligand complex (**B**) or the binary CSE-substrate complex (**C**). As shown, MD simulations induce structural changes as well as modifications of the hydrogen bonding network observed in the docking pose in the presence of D-pen. In particular, the structural elements I and IV becomes connected through a H bond between *Ser56 and Ser358 while the connection between structural elements II and III is disrupted. Importantly, the perturbations recorded in the presence of D-pen during 50 ns MD simulations are not observed in the absence of the ligand, thus suggesting that the fixation of D-pen specifically induces structural changes in the vicinity of the active site and at the interfacial ligand binding site. *Residues from the adjacent subunit B. H-bonding are depicted by light blue lines.

